# UnionLoops: a workflow for calling chromatin loops across related Hi-C datasets with improved specificity, precision, and sensitivity

**DOI:** 10.64898/2026.01.19.700311

**Authors:** Jiangyuan Liu, Johan H. Gibcus, Job Dekker

## Abstract

Chromatin loop calling from chromatin interaction data often exhibits substantial variability across related samples. We present UnionLoops, a computational workflow for chromatin loop calling across multiple related samples. UnionLoops integrates information across datasets to determine positions and dataset-specificity of looping interactions. It constructs a unified candidate loop set, applies consistent filtering and aggregation, and evaluates loop support across samples. We demonstrate that UnionLoops increases sensitivity for detecting shared chromatin loops, reduces spurious sample-specific calls, and improves concordance with independent genomic features, including CTCF and cohesin occupancy. UnionLoops enables improved biological interpretation of chromatin loop organization and dynamics across related conditions.

## Background

Genome folding into a 3D nuclear organization is the result of an interplay between compartmentalization and loop extrusion [1]. Both processes can give rise to localized long-range interaction between defined loci. For instance, microcompartmentalization of small regions enriched in histone H3 lysine 27 acetylation (H3K27Ac) can associate through affinity-driven interactions, a microcompartmentalization phenomenon [2,3]. However, the most dominant process that gives rise to specific looping interactions in interphase cells is loop extrusion by the cohesin complex [4]. Cohesin complexes bind DNA throughout the genome, with a preference for specific cis-elements such as enhancers. Through an active process of extrusion, these complexes can then generate loops of up to hundreds of kilobases. Extrusion can be blocked at specific locations, especially at sites bound by the CTCF protein. Such blocking is dependent on the orientation of the bound CTCF protein. This results in positioned loops at pairs of CTCF sites. Loops can also form at other locations, and distal enhancers may be brought into physical contact by cohesin and/or by microcompartmentalization.

Such positioned loops can be detected by chromosome conformation capture-based methods, such as Hi-C and Micro-C, as well as capture-based approaches, including ChIA-PET and PLAC-seq [5,6]. In datasets obtained with these methods, looping interactions can be discerned as focal enrichments (dots) in interaction matrices representing frequent interactions between pairs of DNA loci called loop anchors. Loop anchors are often pairs of CTCF binding sites in a convergent orientation. These loops are the hallmarks of blocked loop extrusion by cohesin [7] and represent a large fraction of loops that are typically detected in Hi-C and Micro-C data [6]. Although perhaps less prominent, loops can also be detected in other places where active loop extrusion is stalled, including active promoter-enhancer interactions [1]. An increase in such interactions can result in local gene expression changes.

Several algorithms have been developed to qualify and quantify focal interaction directly from interaction maps [7,8]. Widely used approaches include HiCCUPS and SIP. Although both methods work well, they differ quantitatively: HiCCUPS is relatively conservative, by removing weaker loops, i.e., those represented by a single pixel in a Hi-C heatmap. This leads to a low false positive rate. In a direct comparison with HiCCUPS, SIP reduced the number of false negatives, while preserving most true positive loop calls [8]. These methods are designed to call loops in single datasets, but are often used to perform comparative loop calling between datasets [9,10]. However, none of these algorithms were developed specifically to detect differential focal interactions between maps of related samples, e.g., Hi-C datasets representing different time points along a specific treatment of the cells.

With the maturation of chromosome capture techniques, comparisons of genome folding between related samples have become more mainstream. Several recent publications have applied established focal interaction detection algorithms to detect changes in looping in a series of related samples [11,12]. When such analysis is performed with conventional HiCCUPS or SIP, this has been shown to lead to errors in defining loops as dataset-specific or as shared loops [13]. We hypothesize that the detection of focal interaction maps can be improved by aggregating information from related samples. We present UnionLoops, a nextflow pipeline based on HiCCUPS detection that uses the relation between samples. We show that by using information from related samples, UnionLoops significantly improves sensitivity, positional precision, and dataset specificity of focal interaction detection.

## Results

### Applying HiCCUPS to related datasets

HiCCUPS is a widely used method for calling loops in Hi-C data [7]. HiCCUPS involves several steps. First, pixels (pairwise interactions between two bins) that are statistically significantly enriched in the Hi-C signal are identified. Nearby significant pixels are then clustered, and a centroid pixel, i.e., the pixel with the strongest Hi-C signal, representing the cluster, is determined. This centroid pixel then needs to pass additional heuristic thresholds [7]. Additionally, when only a single pixel is significant, with no nearby pixels identified, this would represent a cluster of 1 pixel, i.e., a “singleton”. Singletons need to pass more stringent thresholds to be included in the final loop list. HICCUPS is very conservative, and weak loops, especially those represented by just a single significantly enriched pixel in the Hi-C map, are often removed, which can eliminate true positives. Further, HiCCUPS was not designed to perform comparative loop calling between datasets, but is regularly used for such analysis [9,10]. The potential removal of true positives and the use of conventional HiCCUPS for differential loop calling across datasets can lead to inaccurate results, as we illustrate below (Fig. 1; Additional file 1: Fig. S1a).

**Fig. 1.**
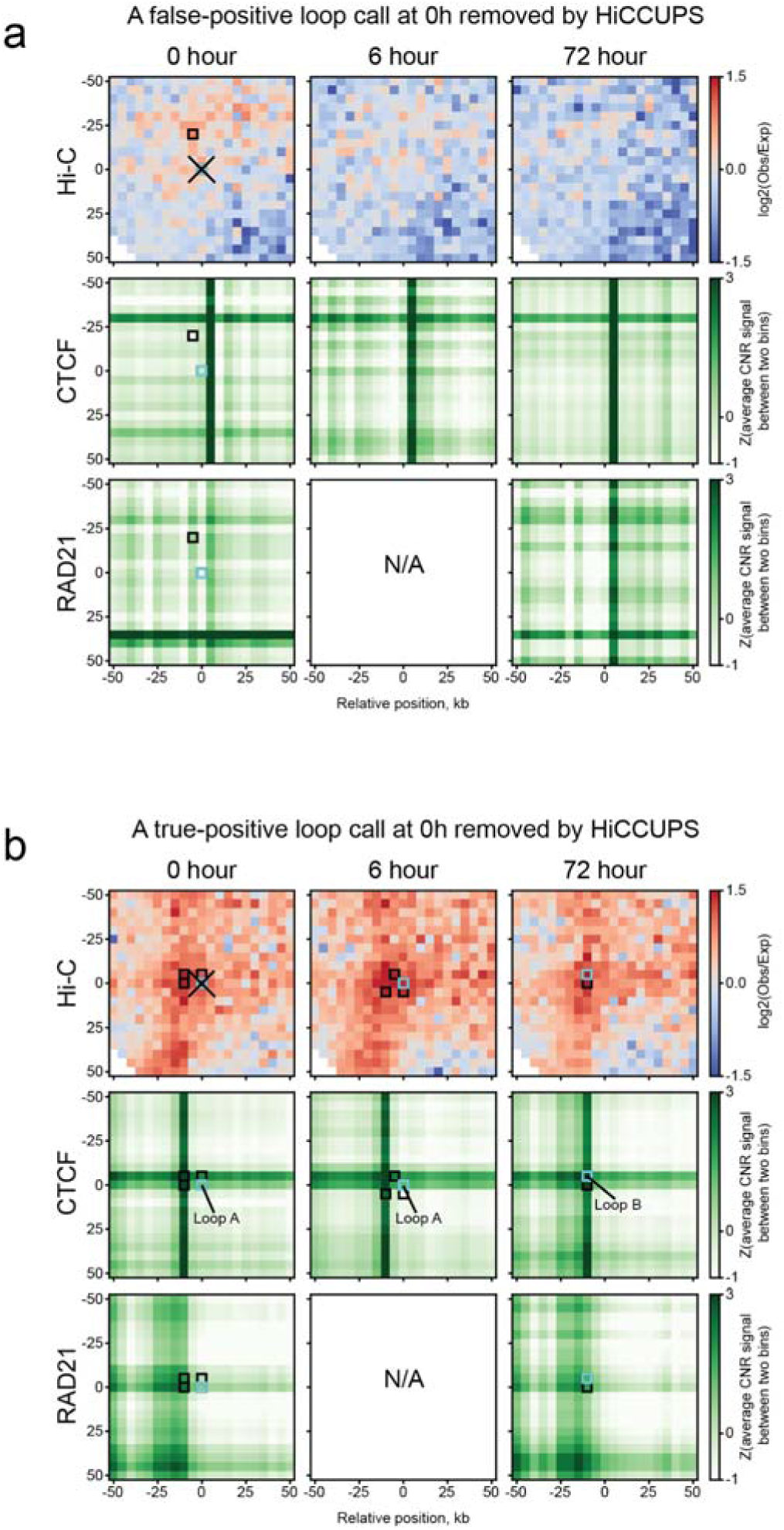
HiCCUPS stringent filtering on called loops. **a**, A removed false positive loop (light blue) at t = 0 h in a cluster of enriched pixels (black) shown in the same snippet of Hi-C maps and element-wise z-score normalized matrices of average Cut&Run signals (read count) of CTCF and RAD21 between 5 kb row–column bin pairs at t = 0, 6, and 72 h (see Methods). No pairs of CTCF/RAD21 binding sites (i.e., crossing points) are adjacent to both anchors of this loop at all three time points. **b**, A removed true positive loop (light blue, loop A) at 0 h and the other two loops (light blue, loop A and B) at 6 and 72 h, respectively, shown in the same way as in **a**. Their clusters of enriched pixels are overlapped with a pair of CTCF/RAD21 binding sites at all three time points. In particular, both anchors of loop B at 72 h are precisely overlapped with a pair of CTCF binding sites. Coordinates of the center pixel: **a**: chr16: 25735000-25740000 & chr16: 25830000-25835000; **b**: chr11: 5205000-5210000 & chr11: 5300000-5305000.

We used HiCCUPS to identify loops on a series of related Hi-C datasets. We first selected the three-time-point in situ Hi-C data from Bond et al., where Hi-C was performed on K562 cells differentiated into a megakaryocyte-like state for 0, 6, and 72 hours [11]. We mapped and processed these Hi-C datasets using the Distiller pipeline (see Methods), resulting in approximately 2.5 billion valid intra-chromosomal reads per timepoint after merging four biological replicates [14]. For each differentiation timepoint we then applied conventional HiCCUPS on 5 kb-resolution Hi-C maps separately and identified 4088, 3030, and 5818 dots representing DNA loops (Additional file 2: Table S1). We next applied the default stringent filtering per time point using the thresholds set by HiCCUPS (see Methods). Bond et al. also generated CUT&RUN datasets for CTCF and RAD21 per time point, which allowed us to validate loops identified by HiCCUPS.

We compared loop calls between the timepoints and overlapped them with RAD21 and CTCF CUT&RUN data from the same study to validate and confirm that the stringent filters imposed by HiCCUPS accurately removed false positive loops. In the example shown in Fig. 1a, at t = 0 hours, HICCUPS identified a cluster of 2 significant pixels. The centroid of this cluster (light blue outlined pixel) did not pass the centroid threshold, and thus the loop was removed. Comparison to CTCF and Rad21 Cut&Run data (lower heatmaps) showed that this pixel did not overlap with binding sites of these proteins, suggesting this was not a true looping interaction, and HICCUPS correctly removed this call from the final loop list. We also note that no significant pixels were identified at t = 6 and 72 hours in this region.

However, HICCUPS also removed true positives (Fig. 1b). In this example, significant pixels were identified in the same section of the Hi-C heatmaps for t = 0, 6, and 72 hours. For each of the three time points, centroids were identified for each cluster of significant pixels. The centroid at t = 0 hours did not pass the centroid filtering, and the loop call was removed. However, we note that at all time points, including t = 0 hours, the called loop involves loop anchors that contain CTCF-bound sites. This strongly suggests that the loop is also present at t = 0 hours, and HICCUPS removed a true positive. In addition, we note that HICCUPS placed the centroid at the same position for t = 0 and 6 hours but at a different location for t = 72 hours, even though the clusters of significant pixels at different timepoints partially overlap. This leads to two different but very closely positioned loop calls for the three datasets. This can lead to the conclusion that there are timepoint-specific loops in this region. Alternatively, the loops are the same, i.e., involving the same pair of loci, in all three datasets; however, there is imprecision in defining the exact location of the loop, leading to three estimates that differ slightly.

Problems in calling loops specific to a given dataset are perhaps best illustrated by calling loops for two replicates for K562 cells at t = 0 hours that should not show any differential looping interactions. We generated a set of two technical replicates by pooling Hi-C replicates 1 and 2, and Hi-C replicates 3 and 4. We then performed the conventional HiCCUPS and found that the largest categories of loops are dataset-specific (Additional file 1: Fig. S1a). In many cases, this arises from imprecision of loop call positions, similar to what we illustrate in Fig. 1b.

To address these and other limitations of loop calling based on Hi-C data, we developed UnionLoops, a workflow for loop calling using a set of related Hi-C datasets. UnionLoops is motivated by two hypotheses: First, we found that whereas true positive loops were often shared across related samples, false-positive loops, likely arising from technical variation, appeared largely sample-specific and rarely overlapped with those in related samples (Fig. 1a-b). We hypothesized that comparing loops among related samples would be effective in rescuing true positives mistakenly discarded due to HiCCUPS’ stringent thresholds. Second, we hypothesized that defining the position of a loop once by identifying the centroid of a cluster of pooled enriched pixels across multiple related samples, rather than independently within each sample, could improve both loop calling accuracy and specificity. Below, we describe UnionLoops and show that it achieves: (1) higher sensitivity by rescuing true positive loops; (2) greater positional precision; and (3) improved loop specificity.

### UnionLoops: a workflow for loop calling across datasets

To investigate chromatin loop dynamics across multiple related samples, i.e., a series of Hi-C datasets representing a time series upon some treatment or across differentiation, etc., it is essential to construct a precise and comprehensive union list of annotated loops with sample-specific information. We compared the conventional method used in two previous studies [9,10], wherein loop specificity was based on exact overlaps of HiCCUPS loops from individual samples, with our new method, UnionLoops, which integrates loop calling and comparison across multiple samples within a unified HiCCUPS-based workflow (Fig. 2).

**Fig. 2.**
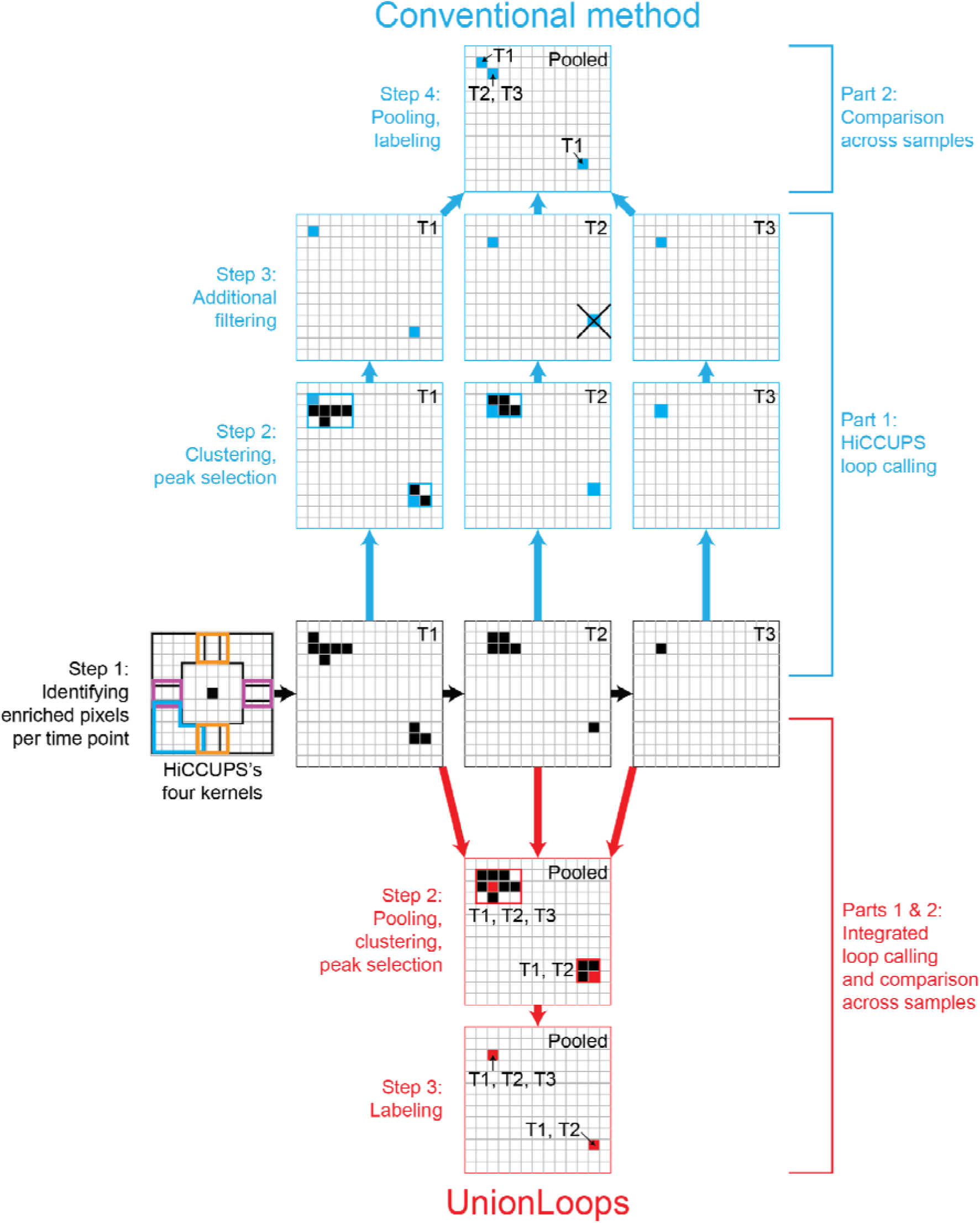
Schematic flowchart illustrating the workflows of the conventional method and UnionLoops to generate a union list of labeled loops across three time points (T1, T2, T3). Grids, snippets of Hi-C maps of different samples at the same genomic loci; black pixels, enriched pixels identified by HiCCUPS; rectangles, clusters of enriched pixels; blue/red pixels, peak pixels in clusters; the cross-marked pixel, a removed loop that does not pass HiCCUPS’s stringent filtering threshold.

The schematic flowchart in Fig. 2 illustrates how the conventional HiCCUPS method and UnionLoops are used to identify loops within the same region from Hi-C maps of three time-series samples (T1, T2, and T3). In the conventional method, HiCCUPS is first used to identify significantly enriched pixels at the three time points separately (Step 1). Secondly, HiCCUPS performs clustering on these enriched pixels for each time point and determines the loop position by identifying the peak pixel (centroid) for each cluster at each time point (Step 2, blue). Then, to minimize the number of false positive loop calls, HiCCUPS performed additional filtering on two types of loops: centroids of clusters of significant pixels needed to pass a centroid-specific threshold, while singletons needed to pass an even more stringent threshold (Step 3, blue). Fourth, for the dataset-specificity analysis, the conventional method pooled the final HiCCUPS loops from each of the three time points and annotated them according to their respective time points (Step 4, blue).

In contrast, after identifying enriched pixels for each time point using HiCCUPS as in the conventional method (Step 1), UnionLoops first pools all significant pixels observed across the three datasets, and then performs the same HiCCUPS-based clustering framework and identification of the centroid pixel once (Step 2, red). In this approach, the centroid is defined as the pixel with the maximum sum of normalized contacts across the samples from which the clustered enriched pixels were derived. The selected peak pixels (centroids, representing the final loop calls) are labeled based on the samples contributing to the enriched pixels in their clusters (Step 3, red). In effect, UnionLoops uses information from multiple datasets to define the position of loops. As we show below, this leads to greater precision in defining the position of the loops.

By pooling significant pixels from different datasets, many significant pixels that were singletons in individual datasets become part of a cluster when nearby pixels are identified in the other datasets. The centroid of such a cluster then defines the final loop call. This is expected to lead to the identification of more loops because the more stringent threshold for singletons does not apply anymore.

For any remaining singletons, UnionLoops applies an additional two-step filtering process to remove (1) those singletons whose anchors did not overlap with boundaries of any non-singleton clusters, i.e., do not share a loop anchor with other called loops (Additional file 1: Fig. S2); and (2) those sample-specific singletons that did not pass the stringent HiCCUPS q-value threshold similar to the conventional HiCCUPS method.

UnionLoops was designed to (1) rescue true positive loops accidentally removed by HiCCUPS at each time point by looking for nearby enriched pixels from other related samples (e.g., the bottom right loop at T2); (2) better define loop specificity by clustering pooled enriched pixels between multiple related samples (e.g., one shared loop across T1, T2, and T3 instead of one T1-specific loop and one T2, T3-specific loop at the upper left); and (3) fine-tune the location of each loop within its cluster using normalized contacts aggregated from all involved samples, rather than from individual samples (e.g., both upper left and the bottom right loops) (Fig. 2).

### Application of UnionLoops to two series of Hi-C datasets

To compare loop-calling performance, we applied both the conventional method and UnionLoops to Hi-C data from two independent studies. We used data from three-time points of in situ Hi-C data from Bond et al., and a five-time-point in situ Hi-C dataset of pancreatic cell differentiation from Lyu et al. [11,12]. The latter study had shallower sequencing depth, averaging ∼650 million valid intra-chromosomal reads per time point, and included Hi-C data obtained with H9-hESCs differentiated into definitive endoderm (DE), primitive gut tube-like (PGT), pancreatic progenitors (PP), and stem cell-derived β-cell organoids (SC-β organoids) [12]. We applied the conventional method and UnionLoops with Hi-C data binned at the same resolution (5kb) and using the same radius (20kb) for clustering enriched pixels per time point [7]. We also generated two “MEGA” Hi-C maps by merging all individual Hi-C maps from each study, combining the three time points from Bond et al. and the five time points from Lyu et al., respectively [11,12]. MEGA Hi-C maps display a higher signal-to-noise ratio and sensitivity, which enables the detection of weaker loops (as shown in Bond et al. and below) [11].

We found that UnionLoops was able to rescue a modest number of additional time point–specific loops by clustering them with nearby enriched pixels uniquely identified in the MEGA map (Fig. 3a; Additional file 1: Fig. S3a-b). Furthermore, UnionLoops fine-tuned their positions (e.g., a 0-hour-specific loop) by selecting the loops with the strongest aggregated signals across relevant Hi-C maps (e.g., 0 hour and MEGA), thereby improving positional precision, as evidenced by enhanced overlap with CTCF/RAD21 binding sites (Fig. 3a). At the same time, even after including the MEGA map as an additional input for pooling in UnionLoops, a noticeable portion of singletons in each Hi-C map remained isolated, i.e., failed to cluster with enriched pixels from any other Hi-C maps, indicating a need for additional singleton filtering as described above (Additional file 1: Fig. S4). Therefore, UnionLoops more moderately used HiCCUPS’s extremely stringent q-value threshold to only perform filtering on these “persistent” singletons instead of all singletons identified at individual samples by HiCCUPS. In contrast, because only a small fraction of centroids remained that did not overlap with any statistically significantly enriched pixels in another dataset in a single Hi-C map, UnionLoops retained them instead of applying HiCCUPS’s local enrichment filtering for centroids (Additional file 1: Fig. S4).

**Fig. 3.**
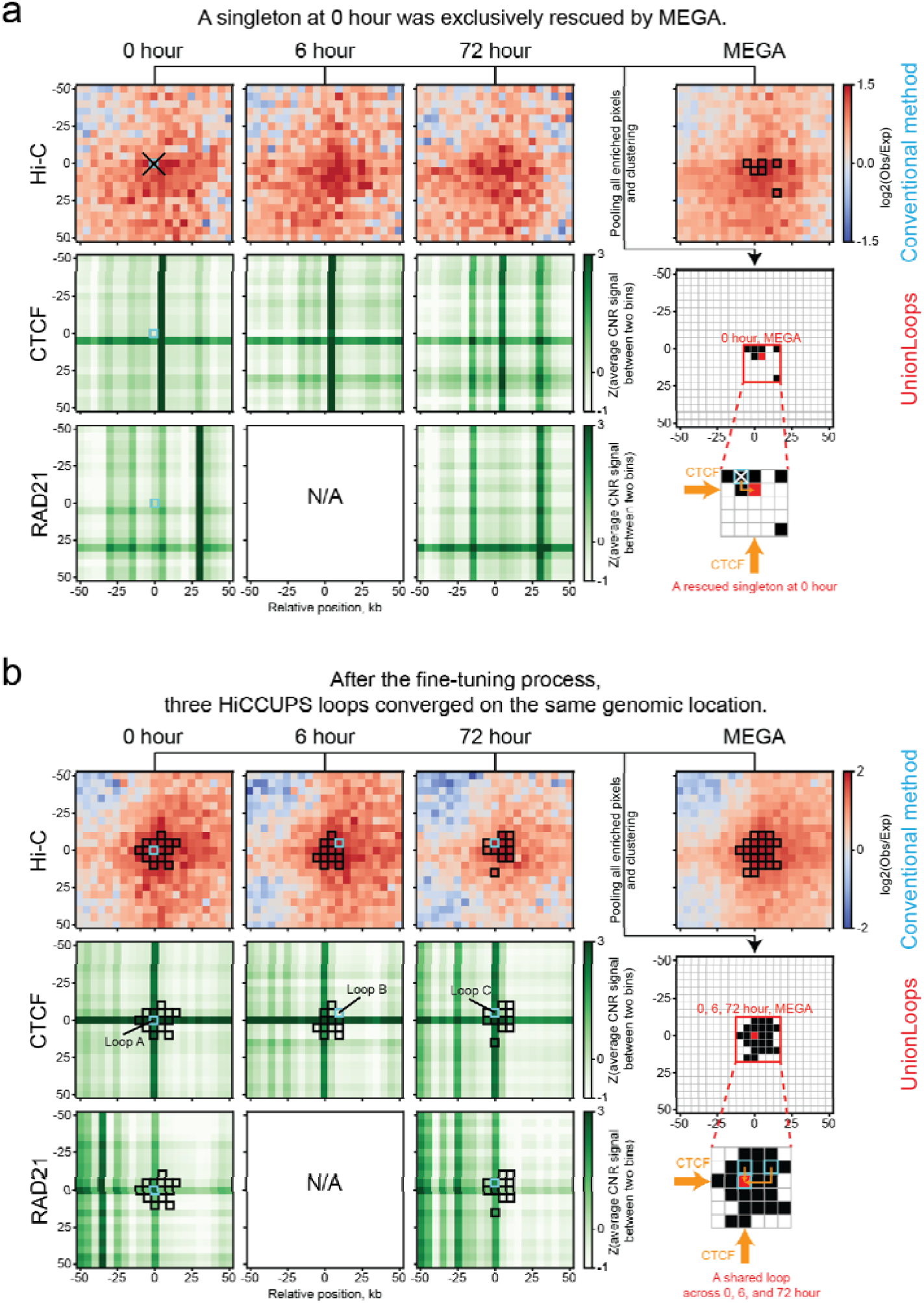
UnionLoops can fine-tune the genomic locations of both rescued and HiCCUPS loops to achieve improved definition of loop specificity. **a**, A true-positive loop (light blue) at 0 h, excluded by the HiCCUPS singleton filter, was rescued as a 0 h-specific loop after fine-tuning its position within the cluster of enriched pixels (black) identified from the 0 h and MEGA Hi-C maps. **b**, Three HiCCUPS loops (light blue) at 0, 6, and 72 hours, respectively, were defined as a shared loop across all three time points after fine-tuning their locations within the same cluster of enriched pixels (black) pooled from Hi-C maps of three time points and MEGA. Matrices of Hi-C, CTCF, and RAD21 are shown in the same way as in Fig. 1. Coordinates of the center pixel: **a**: chr10: 110740000-110745000 & chr10: 110885000-110890000; **b**: chr10: 32795000-32800000 & chr10: 33005000-33010000.

Besides those rescued loops, UnionLoops retained nearly all HiCCUPS loops per sample, except for those singletons whose anchors did not overlap with the boundaries of any non-singleton clusters after pooling all samples, representing only a small fraction of the total HiCCUPS loops per sample (for Bond et al. dataset: 1.7% (0 hour), 1.5% (6 hour), and 2.8% (72 hour); for Lyu et al. dataset: 2.3% (hESC), 1.8% (DE), 1.1% (PGT), 2.4% (PP), and 4.6% (SC-β organoids)) (Additional file 1: Fig. S2). As with the rescued loops, UnionLoops clustered pooled enriched pixels of these HiCCUPS loops from all Hi-C maps, allowing positional fine-tuning across relevant samples within each cluster and aligning them to a shared genomic location rather than permitting slight sample-specific variation in loop position. For example, the conventional method identified loops A, B, and C specifically at 0, 6, and 72 hours, respectively.

However, the anchors of loops B and C were misaligned to a pair of CTCF/RAD21 binding sites shared across all three time points (Fig. 3b). UnionLoops fine-tuned loops A, B, and C to the same position as a shared loop across three time points, which was perfectly aligned to that pair of CTCF/RAD21 binding sites. Therefore, compared to the conventional method, the two examples in Fig. 3 suggest that UnionLoops can fine-tune the positions of both rescued and HiCCUPS loops in each sample, thereby enabling a more accurate definition of loop specificity and positioning across related samples.

### UnionLoops vs. HiCCUPS: improved sensitivity and positional precision

In addition to the two examples of fine-tuned rescued and HiCCUPS loops shown in Fig. 3a and 3b, respectively, we systematically evaluated how UnionLoops improves the sensitivity and precision of loop identification compared to the original HiCCUPS loops used in the conventional method. At each time point, UnionLoops rescued a substantial number of loops, compared to those identified by HiCCUPS, in most cases doubling the number of loops (Fig. 4a; Additional file 1: Fig. S5a). It enhances sensitivity by pooling detection across datasets, allowing recovery of loops that do not meet the filtering threshold in individual runs. Aggregated interaction data revealed that rescued loops consistently exhibited weaker signal strength, leading to their exclusion by HiCCUPS’s stringent thresholds (Fig. 4b; Additional file 1: Fig. S5b).

**Fig. 4.**
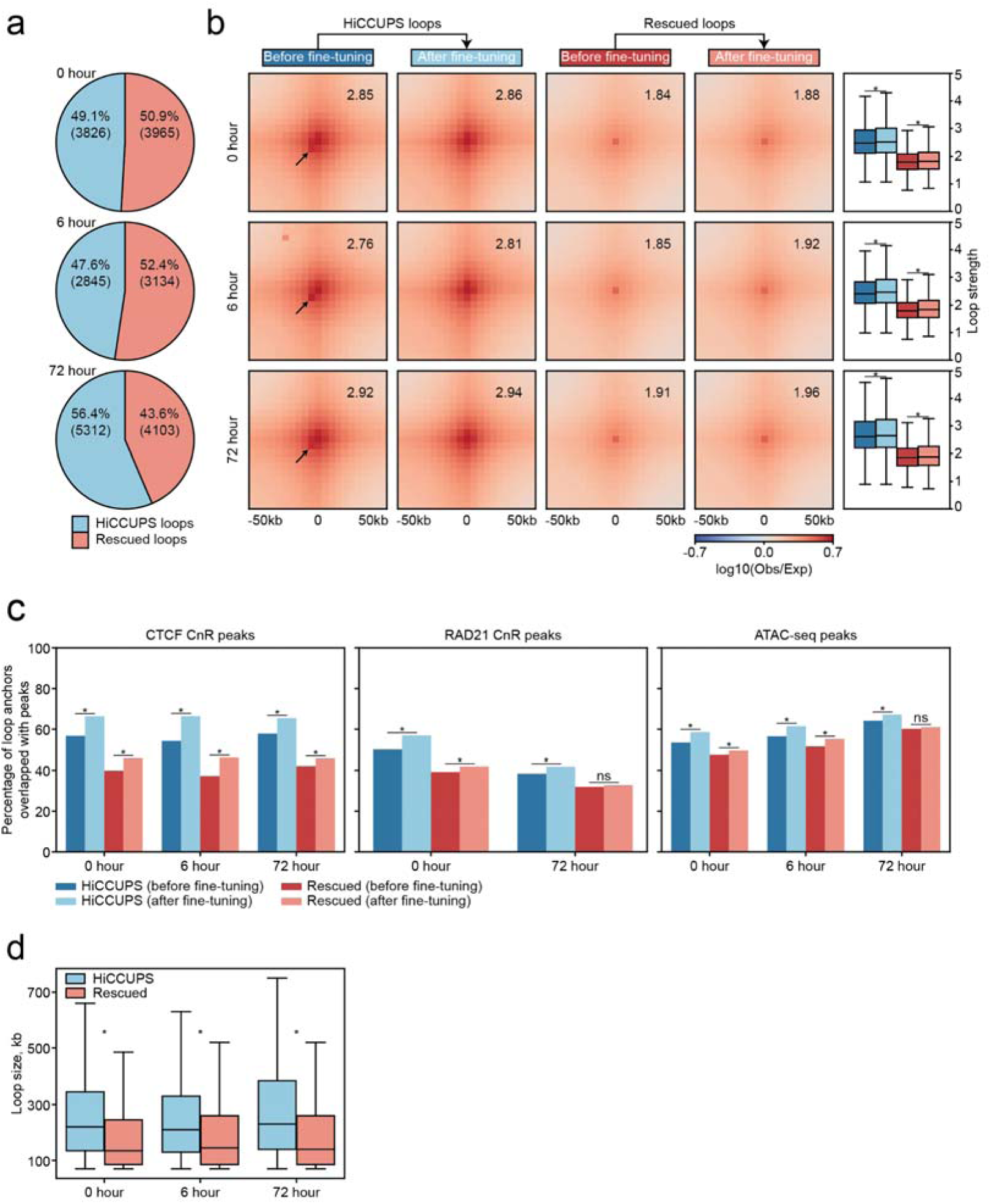
UnionLoops improves sensitivity and positional precision of HiCCUPS loop calls at three time points. **a**, The percentage of HiCCUPS loops versus the percentage of rescued loops. **b**, Pileups of HiCCUPS and rescued loops before and after fine-tuning, with corresponding loop strength distributions. The averaged loop strength value in each pileup represents signal enrichment relative to the local background (see Methods). Arrows highlight focal interactions that deviated from HiCCUPS loops at the center of pileups before fine-tuning. Wilcoxon rank-sum test; asterisks indicate P < 0.05. **c**, The percentages of anchors from HiCCUPS and rescued loops, before and after fine-tuning, that overlapped with CTCF CnR, RAD21 CnR, and ATAC-seq peaks. Chi-square test of independence; asterisks indicate P < 0.05. **d**, Loop size comparisons between HiCCUPS and rescued loops. Wilcoxon rank-sum test; asterisks indicate P < 0.05. Loop resolution is 5 kb for all panels (**a–d**).

In addition to increased sensitivity, UnionLoops further fine-tuned the positions of both HiCCUPS and rescued loops by selecting representative pixels within clusters of pooled enriched signals from related samples. Specifically, we observed that, before fine-tuning, not all HiCCUPS loops were centered on the pixels with the highest interaction frequency within loop pileups, as indicated by the arrows in Fig. 4b. Such deviations were not observed in the five time points reported by Lyu et al., which may be attributed to the lower read depth in their data (Additional file 1: Fig. S5b, and see below). Following fine-tuning by UnionLoops, all HiCCUPS loops were more precisely aligned to the central pixels with the strongest interaction signals, resulting in a more focused and radially symmetric aggregated Hi-C signal pattern at loops across all three time points. This improved positioning of both HiCCUPS and rescued loops was accompanied by increased loop strength (Wilcoxon rank-sum test; P < 0.05), as an additional output from our UnionLoops nextflow pipeline for quantitative downstream analyses (Additional file 1: Fig. S7) [15]. These results provide further evidence for enhanced positional precision.

Beyond loop strength, we also evaluated the improvement in accuracy of fine-tuned loops by assessing whether their anchors overlapped with CTCF/RAD21 binding sites and chromatin accessibility sites defined by ATAC-seq peaks (Fig. 4c; Additional file 1: Fig. S5c). After fine-tuning by UnionLoops, both HiCCUPS and rescued loops exhibited significantly higher percentages (Chi-square test of independence; P < 0.05) of anchors overlapping with CTCF/RAD21 binding sites and chromatin accessibility sites at each time point in both the Bond et al. and Lyu et al. datasets. This particularly supports that the rescued loops, despite their weak loop strength, were true positives inadvertently excluded by HiCCUPS’s stringent thresholds. We also found that these rescued loops with weak loop strength tend to be smaller in genomic distance compared to the HiCCUPS loops with strong loop strength in the Bond et al. dataset (Wilcoxon rank-sum test; P < 0.05) (Fig. 4d). This is consistent with the notion that short-range loops are more difficult to detect due to the high background contact frequency in Hi-C data at smaller genomic distances. This challenge likely explains why HiCCUPS applies stringent filters, sacrificing sensitivity to maintain low false positive rates. We observed that, unlike the Bond et al. dataset, HiCCUPS loops are not larger than rescued loops at all time points (i.e., hESC, DE, and PGT) in the Lyu et al. dataset, which might be due to the lower sequencing depth in the Lyu et al. dataset (Additional file 1: Fig. S5d, and see below). Therefore, after fine-tuning by UnionLoops, both HiCCUPS and rescued short-range loops were more precise and more accurate, as evidenced by stronger loop strength and improved overlap with CTCF/RAD21 binding sites and chromatin accessibility sites.

We note that both loop strength and overlap with CTCF/Rad21 of the set of rescued loops are lower than for the set of fine tuned loops. While their lower interaction strength can explain why they were filtered out by conventional HiCCUPS, we can currently not rule out that by rescuing this set of loops and thereby increasing sensitivity, we inadvertently increased the false positive rate. In the absence of a gold standard list of loops we can currently not resolve this issue. That said, the fact that all rescued loops are detected in more than one dataset would support their inclusion.

### UnionLoops vs. the conventional method: improved loop specificity

Next, we systematically evaluated how UnionLoops affects loop specificity, i.e., whether a loop occurs only in a specific sample or is shared between samples. We compared the number of loops at 5 kb resolution detected in one or more samples using the conventional method versus UnionLoops. Application of the conventional method yields primarily time point-specific loops. UnionLoops, in contrast, revealed that the largest set of loops is shared across all time points in the Bond et al. dataset (Fig. 5a). This discrepancy stems from the conventional method’s direct summation of all HiCCUPS loop calls, which artificially makes loops appear unique to each time point due to the imprecision of the independent calls. In the Lyu et al. dataset, UnionLoops also identified a substantially larger set of shared loops across all time points compared to the conventional method. However, this was only detected when the loop resolution was decreased from 5kb to 10 kb (Additional file 1: Fig. S6a-b). The need to use a lower resolution is attributed to the lower read depth in the Lyu et al. dataset, which reduces the sensitivity of HiCCUPS to detect loops consistently present across all time points (see below).

**Fig. 5.**
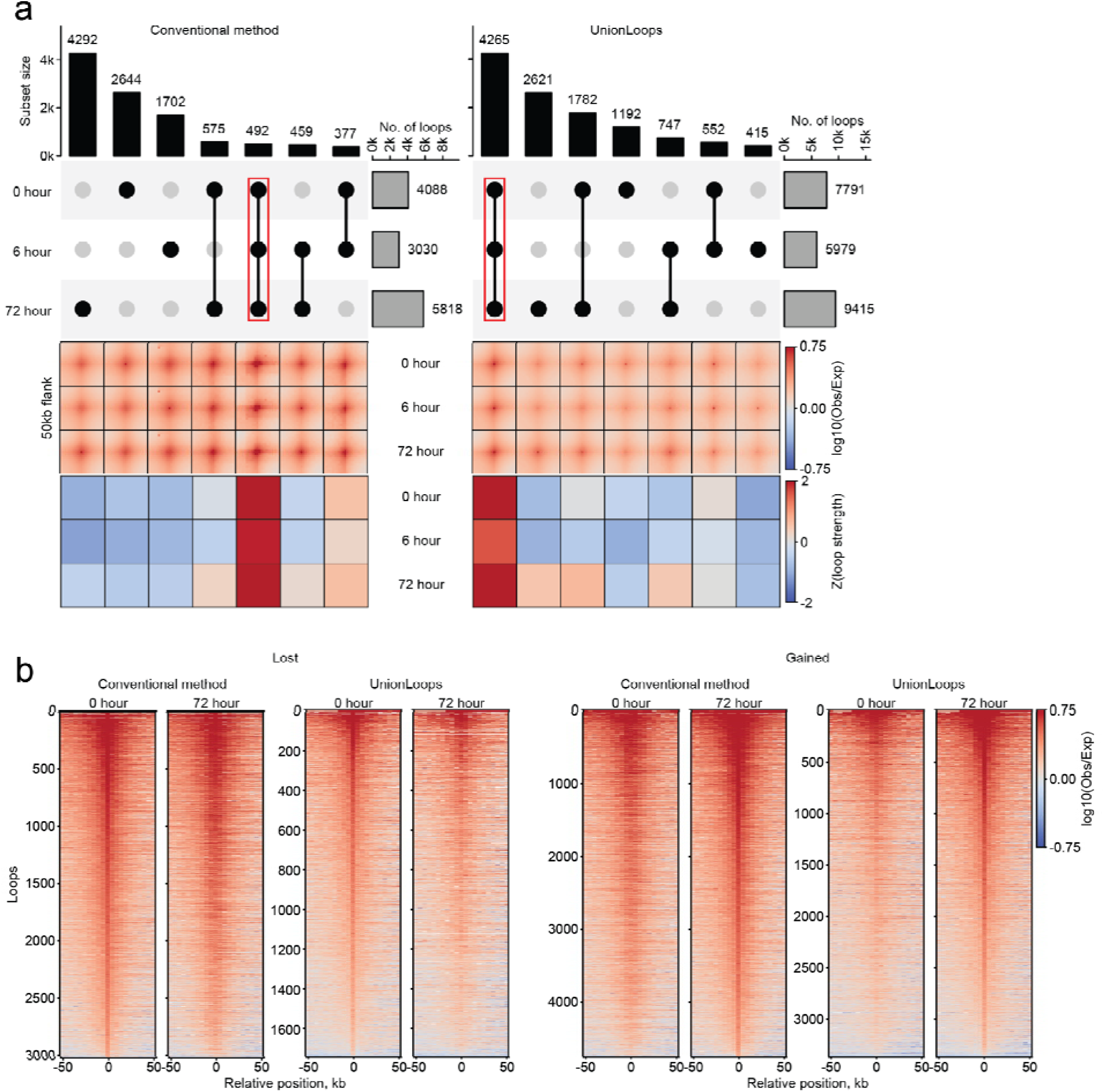
UnionLoops improves loop specificity across 0, 6, and 72 hours compared to the conventional method. **a**, Combined visualization of loop detection and signal strength across the three time points. UpSet plots (top) show the number of loops detected uniquely or in combination across the three time points. Gray bars represent the total number of loops per time point; black bars and connected dots indicate loops detected in one, two, or all three time points; red rectangles highlight loops detected in all time points. Pileups (middle) display averaged contact signals for each loop category defined above. The element-wise z-score normalized heatmaps of the loop strength matrices (bottom) represent signal enrichment relative to the local background illustrated in the pileups (see Methods). **b**, Stackup plots at 50kb-flanked upstream anchors of individual lost and gained loops between 0 and 72 hours. Loop resolution is 5 kb for all panels (**a–b**).

To further validate this finding, we computed the average contact signal and corresponding loop strength for each subset at every time point. The largest subsets of time point–specific loops identified by the conventional method exhibited similar contact signals across all time points, suggesting that these loops are not truly time point specific. UnionLoops demonstrated markedly stronger concordance between loop specificity and contact signal dynamics compared to the conventional method (Fig. 5a; Additional file 1: Fig. S6b).

To address the specificity of loop calls in another way, we examined individual loop contact signals for loops classified as gained or lost by the conventional method in pairwise sample comparisons (Fig. 5b; Additional file 1: Fig. S6c). These loops likewise showed comparable contact signals in both samples, indicating that imprecise loop calling by HiCCUPS leads the conventional method to overestimate sample-specific loops by the direct combination of all loop calls from different time points. In contrast, UnionLoops better captures loops gained or lost between timepoints, with contact signals that are visibly stronger or weaker, respectively.

### Robust improvements in sensitivity, positional precision, and specificity by UnionLoops across various sequencing depths

We evaluated the performance of UnionLoops across a range of sequencing depths by systematically downsampling the Bond et al. datasets from a ∼2,500 million full dataset to 2,000, 1,500, 1,000, and 500 million reads. We then identified loops using conventional HiCCUPS and UnionLoops at 5 kb resolution (Fig. 6). We observed that the number of loops detected by both HiCCUPS and UnionLoops of each time point follow a saturation curve, where the number of identified loops increases rapidly at shallow sequencing depths but plateaus beyond ∼1,500 million valid intra-chromosomal reads (Fig. 6a). This behavior reflects the exhaustion of gaining detectable high-confidence loops when increasing the number of reads under HiCCUPS’s stringent filtering thresholds. In contrast, the number of loops rescued by UnionLoops exhibits a linear relationship with sequencing depth, indicating that UnionLoops effectively leverages increased sequencing depth to increasingly rescue additional loops. Across all read numbers, the number of rescued loops remains comparable to the number of HiCCUPS loops, doubling the number of total called loops, demonstrating improved sensitivity of UnionLoops as compared to conventional HiCCUPS across a wide range of sequencing depths.

**Fig. 6.**
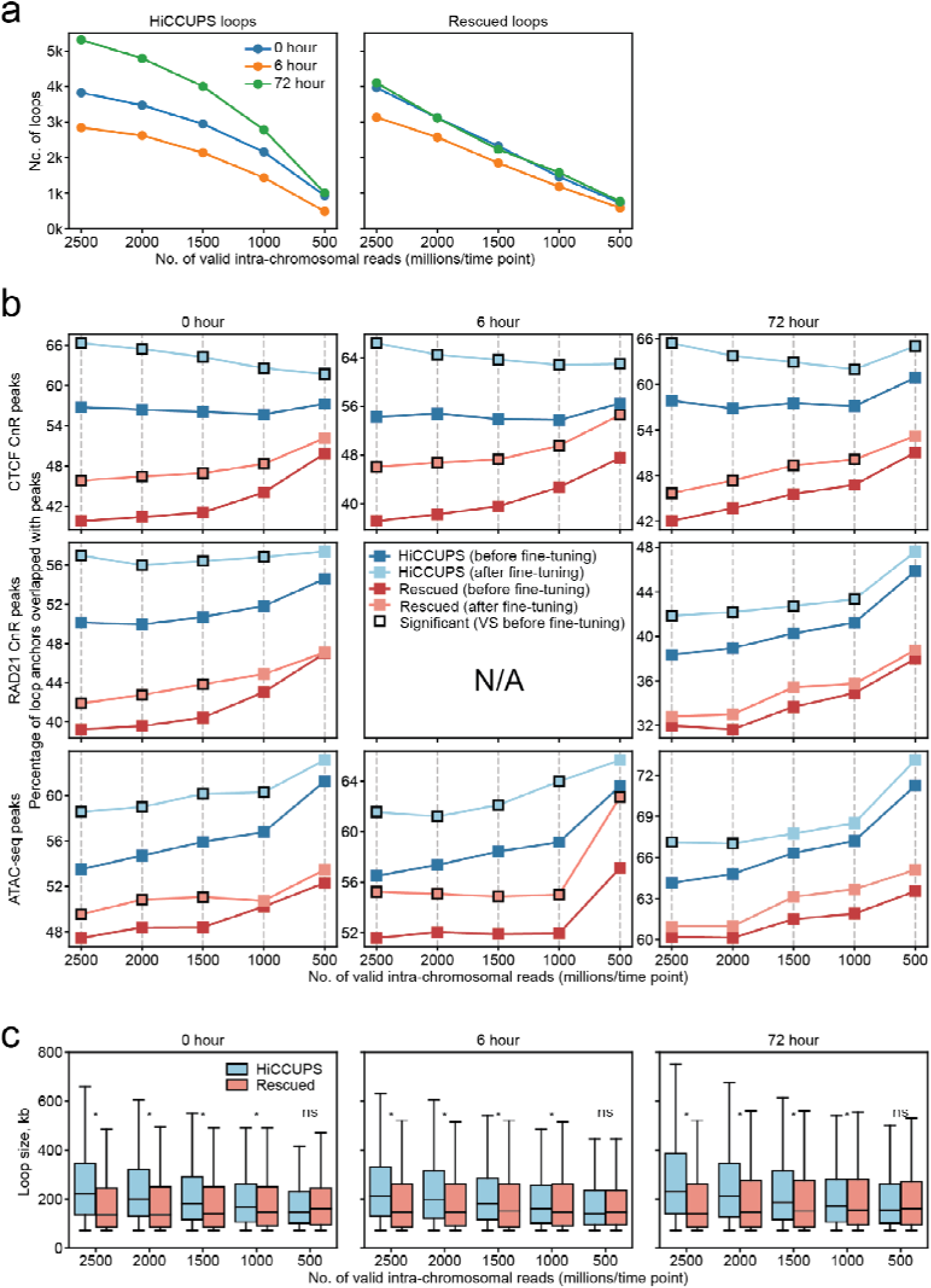
UnionLoops improves the sensitivity and positional precision of HiCCUPS loop calling across various sequencing depths. **a**, Total number of loops detected by HiCCUPS and additional loops rescued by UnionLoops at each time point. **b**, Percentages of HiCCUPS and rescued loop anchors overlapping CTCF CnR, RAD21 CnR, and ATAC-seq peaks before and after fine-tuning across sequencing depths. Y-axis scales differ to better visualize similar trends across different time points. Chi-square test of independence; asterisks indicate P < 0.05. **c**, Loop size comparisons between HiCCUPS and rescued loops across sequencing depths. Wilcoxon test; asterisks indicate P < 0.05. Loop resolution is 5 kb for all panels (**a–c**).

We also assessed whether UnionLoops enhances positional precision of loop anchors at reduced sequencing depths by evaluating their overlap with CTCF, RAD21 binding sites, and ATAC-seq peaks, exactly as described above (Fig. 4c). In this analysis, we compared loop anchors called by conventional HiCCUPS to the same anchors after fine-tuning by UnionLoops. We also compared loop anchors for the set of loops rescued by UnionLoops, before and after fine-tuning. Interestingly, HiCCUPS loop precision increased as sequencing depth decreased, particularly below approximately 1 billion valid intra-chromosomal reads (Fig. 6b). As can be seen from the overlap with Rad21 and ATAC-seq peaks, which increases at lower read numbers. However, this gain in precision came at the expense of sensitivity, with a substantial reduction in the total number of loops detected (Fig. 6a). One interpretation is that at lower sequencing depth, only the strongest loops are detected, and these are called more precisely at Rad21-bound sites. After fine-tuning, UnionLoops further improved the precision of these already strong and high-confidence HiCCUPS loops (Chi-square test of independence; P < 0.05), even at sequencing depths as low as 500 million valid intra-chromosomal reads (Fig. 6b; compare light blue data with dark blue data). Besides rescuing weak loops that conventional HiCCUPS filtered out, UnionLoops also fine-tuned their genomic locations (Fig. 6b; compare light red with dark red data). After fine-tuning, these rescued loops displayed increased overlap with CTCF, Rad21 binding, and ATAC-seq peaks significantly across all read depths (Chi-square test of independence, P < 0.05). The increased overlap of these rescued loops with known regulatory features provides support for their biological relevance, despite the fact that these interactions are relatively weak. Together, these findings demonstrate that UnionLoops effectively mitigates the trade-off between the number of detected loops and their positional precision in HiCCUPS loop calling.

While both the number of detected HiCCUPS loops and rescued loops increased with depth (Fig. 6a), the HiCCUPS loops became progressively biased toward longer-range interactions, whereas the rescued loops identified by UnionLoops remained predominantly shorter-range (Fig. 6c). This pattern likely arises because contact frequency generally declines with genomic distance, making longer-range interactions more difficult to detect at lower depths. In contrast, shorter-range loops are less likely to pass HiCCUPS filters due to high background signals across sequencing depths. These trends help explain the apparent discrepancies in loop size distributions between the deeply sequenced Bond et al. dataset and the shallower Lyu et al. dataset (Fig. 4d; Additional file 1: Fig. S5d). Together, these results demonstrate that UnionLoops primarily rescues shorter-range loops across a range of sequencing depths.

We finally tested whether sequencing depth affects loop specificity across related samples for both the conventional HICCUPS method and UnionLoops. Conventional methods consistently reported more time-point specific loops, regardless of sequencing depth or resolution (Fig. 7-left). In contrast, the largest subset of loops detected by UnionLoops consistently represented loops that were shared across all time points, except at very low sequencing depth (i.e., 500 million valid intra-chromosomal reads) (Fig. 7-right). When we decreased the resolution from 5 kb to 10 kb to increase read coverage per bin, the largest subset of loops detected by UnionLoops was again the set of loops shared across timepoints, while conventional HiCCUPS still detected larger sets of dataset-specific loops. We also observed this phenomenon in the Lyu et al. dataset (Additional file 1: Fig. S6a-b). This observation provides a practical way to assess whether a dataset has sufficient sequencing depth to define loop specificity at a given resolution. These results demonstrate that UnionLoops reliably detects loops with high specificity across a broad range of sequencing depths, and point out where lower resolution may be necessary when depth is limited.

**Fig. 7.**
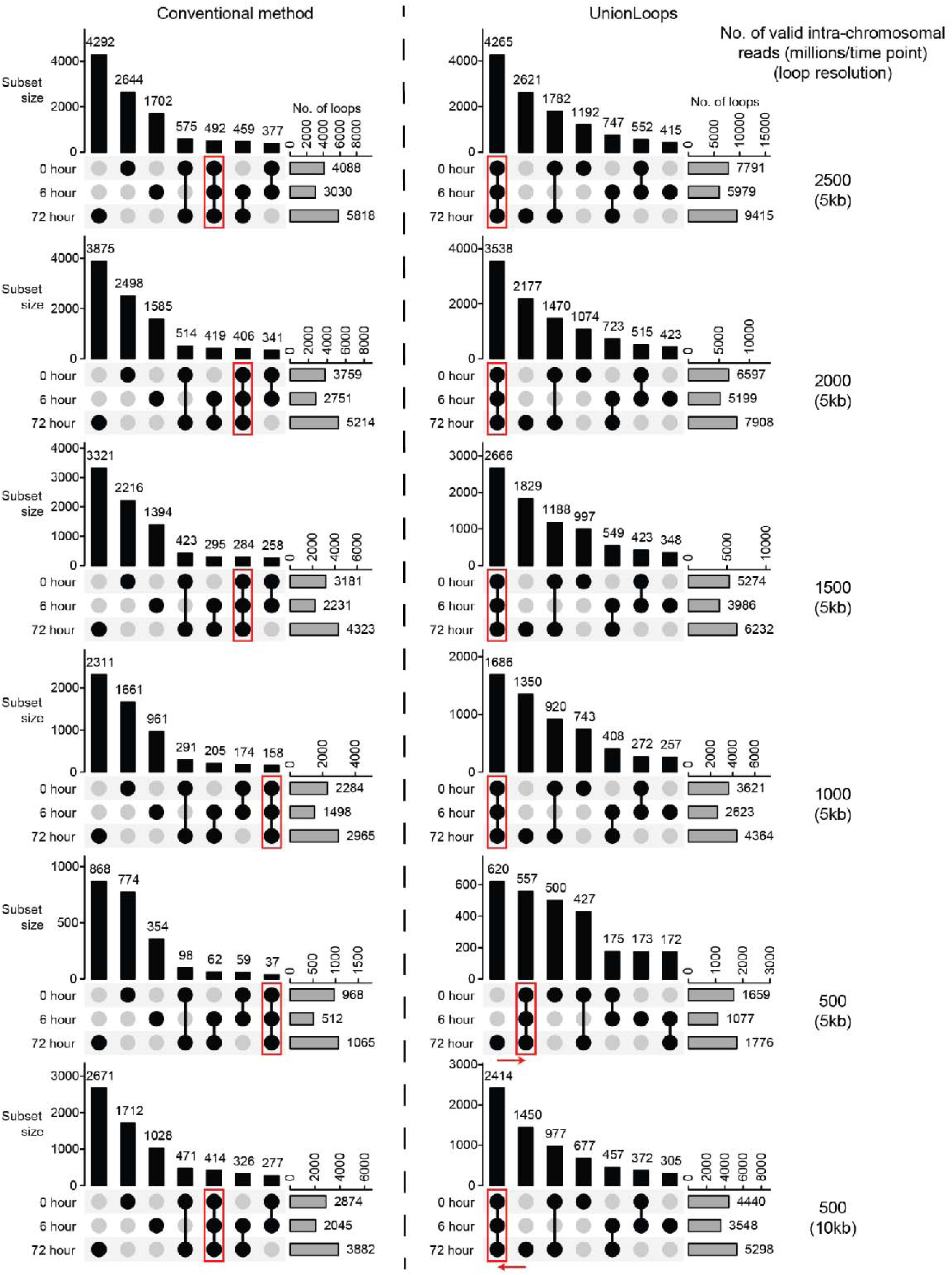
Compared to the conventional method, UnionLoops improves loop specificity across various sequencing depths. UpSet plots are shown in the same way as in Fig. 5a. Red arrows highlight how sequencing depth and loop resolution affect the number of loops detected at all time points by UnionLoops.

### Comparison to related approaches

Others have also realized that loop calling can be imprecise, leading to errors in calling dataset-specific loops. For instance, to address this issue, the Mariner R package performs multi-sample clustering and representative loop selection, similar in concept to our UnionLoops approach but with two key differences (Additional file 1: Fig. S8a) [13]. First, Mariner typically clusters final filtered loops, i.e., single pixel final loop calls for each dataset, whereas UnionLoops clusters pre-filtered significant pixels, i.e., including all statistically significant pixels in a given cluster in a single dataset, enabling UnionLoops, but not Mariner, to rescue true-positive loops removed during per-sample filtering. At the same time, because Mariner uses only filtered loops as input, it is restricted to a much smaller candidate set of pixels, giving it fewer opportunities than UnionLoops to find an optimal representative. Second, Mariner selects the representative loop based on the maximum signal in any single sample, making it sensitive to outliers, whereas UnionLoops uses the summed signal across balanced samples, prioritizing consistent multi-sample support. Thus, although Mariner improves the precision of loop calls over raw loop calls, UnionLoops achieves higher precision by leveraging shared signal and a richer multi-sample candidate pool (Additional file 1: Fig. S8b-c).

To assess whether multi-sample clustering improves loop precision beyond HiCCUPS, we applied both Mariner (as described in Bond et al.) and UnionLoops to merge final loop calls from SIP [8]. While both approaches improved precision relative to raw SIP loops, UnionLoops achieved significantly higher precision than Mariner (Chi-square test of independence, *P* < 0.05) (Additional file 1: Fig. S9a). Consistently, applying the UnionLoops to Mustache-derived loops also resulted in a significant improvement in precision (Chi-square test of independence, *P* < 0.05) (Additional file 1: Fig. S9b) [16]. In addition, UnionLoops showed stronger concordance between loop specificity and contact signal dynamics than the conventional method for both SIP- and Mustache-derived loops, consistent with observations for HiCCUPS (Additional file 1: Fig. S10a-b). Together, these results indicate that our UnionLoops nextflow pipeline generalizes across different loop callers to improve loop precision and specificity across datasets.

## Discussion

Here we describe UnionLoops, a workflow for loop calling across Hi-C datasets. A major advance of UnionLoops is its ability to integrate loop evidence across related samples, overcoming a core limitation of methods like HiCCUPS, which are designed to call loops on one sample at a time. By consolidating spatially overlapping loop signals across these datasets, UnionLoops produces more precise and biologically consistent loop sets while enabling reliable multi-sample comparison.

Compared with conventional loop-calling methods that analyze related datasets separately, UnionLoops demonstrates three key improvements: increased sensitivity, precision, and specificity. All three improvements stem directly from UnionLoops’ integrative multi-sample loop calling strategy. First, sensitivity increases because many weak loops that might be attributed to noise in single datasets can be rescued when they consistently appear across related samples. Second, precision improves as multi-sample evidence fine-tunes loop positions, resulting in better overlap of loop anchors with CTCF and RAD21-bound sites. Third, loop specificity becomes more reliable because this integrative approach allows us to clearly distinguish shared loops that are consistently supported across samples from those that are truly sample-specific.

Determining whether a loop is shared across samples or is sample-specific remains a major analytical challenge in 3D genome studies. Often, in what we have referred to as the conventional method, studies have relied on exact anchor-to-anchor coordinate overlap to define shared loops [9,10]. However, noise and technical variability in individual Hi-C datasets can result in slight shifts in loop coordinates, causing biologically identical interactions to be misclassified as sample-specific interactions. UnionLoops overcomes this limitation by integrating and averaging estimates from loop positions from multiple datasets, which leads to fine-tuning of loop positions. We find that this results in loops that better overlap CTCF/RAD21 sites and are more reliably identified as shared between datasets.

Following Bond et al., we also applied DESeq2 to the UnionLoops-defined gained and lost loops using unnormalized pixel counts [17]. These results show that DESeq2 is markedly more conservative in detecting gained or lost loops as compared to UnionLoops (Additional file 2: Table S2). This limitation likely reflects the sparsity of Hi-C pixel counts, which depends strongly on resolution and sequencing depth, as well as limited biological replication, both of which lower the statistical power of DESeq2 to detect gained and lost loops [11,18]. Thus, the conventional method is the most permissive in calling dataset-specific loops, UnionLoops provides a more balanced intermediate approach, and DESeq2 remains the most conservative.

While the largest set of loops UnionLoops identified is shared between datasets, we found that these shared loops can vary considerably in strength. Beyond reporting loop presence or absence, UnionLoops quantifies loop strength, uncovering strength dynamics even within the largest set of loops shared across all samples and enabling more granular analyses of chromatin looping, such as regression analyses correlating loop strength with other chromatin features (Additional file 1: Fig. S7; Additional file 1: Fig. S11).

Based on these considerations we believe that it is preferable to refrain from calling loops present or absent in any given dataset. Instead, one should define a union set of loop positions across datasets, e.g., using UnionLoops, and then perform a statistical analysis for each loop position to determine how loop strength quantitatively varies (or not) across datasets.

Overall, UnionLoop calls are more biologically meaningful, as compared to conventional HiCCUPS calls, in several ways. First, UnionLoops revealed that most loops are shared across samples (Fig. 5a; Additional file 1: Fig. S6b), indicating a more conserved interaction landscape than previously suggested [9,10]. Such conserved architecture may extend to other systems or experiments. Conversely, the improved definition of the subset of loops that are dataset-specific will aid subsequent studies into the factors that determine such cell-state-specific looping. Second, the improved precision of loop anchor definition provided by UnionLoops allows better correlation of looping with specific DNA-binding elements and DNA-binding factors. Finally, the increased sensitivity allows detection of weaker, i.e., less frequent, looping interactions.

We envision that UnionLoops will be applicable to a wide range of studies that collect a series of related chromatin interaction datasets. For instance, experiments where chromatin interactions are detected as a function of time during a biological process, i.e., cells responding to signals, cells differentiating, or progressing through the cell cycle. Other examples include experiments where specific perturbations are induced, e.g., the acute depletion of specific chromatin regulators, and the response of genome folding is determined.

UnionLoops, unlike DESeq2, benefits more from sequencing depth than from biological replicates. Low-depth replicates may introduce non-shared, unreliable loops due to noise or technical variability at the replicate level, which could in turn affect the quality of multi-sample integration, so merging replicates to increase depth is often preferable. That said, UnionLoops can be used to call loops across individual replicates (Additional file 1: Fig. S1b). Both deeper sequencing and more replicates will make loop calling more precise and sensitive. Further, the experimental protocol will affect these considerations also: for instance, Hi-C3.0 and Micro-C have stronger loop signals than earlier versions of the Hi-C protocol [15]. In addition, different loop callers used with UnionLoops have different thresholds, further complicating the assessment of how read depth and replicate numbers affect final loop calls. These issues need to be explored deeper in future studies.

UnionLoops should be applicable for a variety of datatypes, including Micro-C, Capture Hi-C/Micro-C, and pulldown-based chromatin interaction assays (ChIA-PET, HiChIP), and as we show a variety of methods that nominate significant pixels in interaction maps (HiCCUPS, SIP, Mustache).

While UnionLoops improves performance of multiple loop callers, there are several limitations to the approach. In the absence of a gold standard list of loops, false positive and false negative rates cannot be quantified. For the set of loops that do not overlap CTCF sites we have not been able to ascertain whether UnionLoops improved their precision and sensitivity. We note that this is true for all loop calling approaches. UnionLoops in its current implementation can only rescue HiCCUPS loop calls. The reason is that other loop callers such as SIP have already implemented their own statistical thresholds that are less stringent than HiCCUPs, and thus for those the need for rescuing less frequently interacting loops is reduced. We have not applied UnionLoops to highly diverged cell types, where more cell type specific loops can be expected. Regarding implementation, UnionLoops currently only accepts cooler files. We note that other widely used Hi-C file formats (e.g., .hic file) can be readily converted into cooler files with publicly available tools [19,20]. Finally, UnionLoops is currently only available as a nextflow pipeline.

## Conclusions

UnionLoops is a workflow for calling chromatin loops across samples and provides improved precision, sensitivity, and specificity. The output is a union list of loop calls across all datasets that can then be quantitatively analyzed for detection of changes in looping frequency. UnionLoops can be used in conjunction with several loop calling methods such as the widely used HiCCUPS, SIP, and Mustache, and should be applicable to chromatin interaction datasets generated with any of the 3C-based approaches including Hi-C and Micro-C. UnionLoops is freely available as a NextFlow pipeline.

## Methods

### Data processing

#### In situ Hi-C data

The distiller-nf nextflow pipeline (v0.3.3; https://github.com/mirnylab/distiller-nf) was used to process raw FASTQ files for each time point in both Bond et al. and Lyu et al. Hi-C datasets [14]. The complete processing workflow, along with all parameter settings, has been detailed in a previous study [15]. This pipeline ultimately produced multi-resolution cooler (mcool) files for each time point, containing both raw and iteratively corrected contact matrices generated using the iterative correction procedure described in Imakaev et al. [21]. MEGA mcool files of these two Hi-C datasets were created using raw fastq files of all time points. Due to the large sizes of raw fastq files for K562 Hi-C data, we ran the distiller pipeline to generate every mcool file and then delete its fastq files of each biological replicate per time point one after another, which were further merged to a 1kb-resolution cooler file per time point using hictk merge (v1.0.0) [20]. Same as the last step of the distiller pipeline, normalized mcool files of three time points were created using cooler zoomify (v0.9.3). The MEGA mcool file of the K562 Hi-C dataset was created using hictk merge and cooler zoomify for all biological replicates of all three time points. For the downsampling analysis of the K562 Hi-C data in Fig. 6 and Fig. 7, we used cooltools random-sample (v0.5.2) with the --cis-count parameter to randomly subsample cis contacts to four target depths per time point (2000, 1500, 1000, and 500 million reads) [22]. The resulting downsampled 1 kb-resolution cooler files were subsequently converted into normalized multi-resolution mcool files using cooler zoomify [23]. Corresponding downsampled MEGA mcool files were created, exactly as described above.

#### CUT&RUN, ChIP-seq, and ATAC-seq data

We created ATAC-seq peak lists of H9 hESC from M. Snyder’s laboratory and DE, PGT, PP, and SC-β organoids from V. Corces’s laboratory in the same way as was done in the study of K562 differentiation [24]. In brief, we mapped sequencing reads to the hg38 genome using bwa mem (v0.7.17) and removed duplicate reads using PicardTools (v3.3.0, https://broadinstitute.github.io/picard/) [25]. After merging and indexing biological replicates using SAMtools (v1.3.1), we used MACS2 (v2.2.9.1) with the parameters ‘-f BAM -g hs --nomodel --shift -100 --extsize 200 -q 0.01 --keep-dup all -B --SPMR’ to call ATAC-seq peaks on the merged files [26,27].

### Data analysis

#### Conventional method

##### Part 1: Loop detection per time point using HiCCUPS

The cooltools.dots function, a Python reimplementation of HiCCUPS, was used to detect chromatin loops from the mcool file of each time point and MEGA separately. For the Bond et al. dataset, loops were called at 5 kb resolution; for the Lyu et al. dataset, both 5 kb and 10 kb resolutions were used. The following parameters were applied in all cases: lambda_bin_fdr=0.1, max_loci_separation=10000000, tile_size=5000000, max_nans_tolerated=4, clustering_radius=20000, and cluster_filtering=None. Setting cluster_filtering=None invoked HiCCUPS’s empirical filtering strategy to reduce false-positive loop calls.

##### Part 2: Annotation of loop specificity

Loops detected at each sample were merged into a union list by requiring exact coordinate matches for both loop anchors. Each loop in the union list was then annotated with a specificity label indicating its presence or absence across time points.

### UnionLoops

We used the UnionLoops nextflow pipeline (v1.0.0) to generate a union list of chromatin loops with specificity labels, utilizing mcool files from all individual time points and the corresponding merged MEGA library. As shown in Fig. 2, UnionLoops applied the built-in HiCCUPS to identify enriched pixels from each input mcool file, which were then pooled and clustered using the same HiCCUPS-based framework. Within each cluster, a representative pixel was selected as the locus with the maximum summed normalized contact across all datasets. Singleton clusters (i.e., clusters containing a single pixel) lacked adjacent enriched pixels in other datasets and thus did not benefit from UnionLoops; these were retained only if they shared at least one anchor with a non-singleton cluster (Additional file 1: Fig. S2) and satisfied stringent HiCCUPS filtering thresholds specific to singletons. For the Bond et al. dataset, the union list was generated at 5 kb resolution; for the Lyu et al. dataset, union lists were generated at both 5 kb and 10 kb resolutions. The following parameters were applied in all cases: assembly_name=’hg38’, lambda_bin_fdr=0.1, max_loci_separation=10000000, tile_size=5000000, max_nans_tolerated=4, and dots_clustering_radius=20000. For the Bond et al. dataset, resolution=5000 and flank=50000 were used; for the Lyu et al. dataset, resolution=5000 and 10000 with flank=50000 and 100000, respectively, were used.

### Average bigWig signal matrices for CTCF and RAD21 across pairs of interacting genomic bins

Using the pybbi package (v0.3.0), we applied the bbi.stackup function (summary=’mean’) to calculate bigWig signal values from CTCF and RAD21 CUT&RUN data in the study of K562 differentiation [28]. Specifically, this function took two vectors of 5 kb genomic bins, corresponding to the rows and columns of the respective Hi-C map snippet, and extracted their average bigWig signals across all time points. We then computed the outer sum of these two vectors of bigWig signal values and divided the result by two to obtain their pairwise averages. Finally, the resulting matrices were subjected to element-wise z-score normalization to enable direct comparison across multiple samples (as in Figs. 1a–b and 3a–b).

### Evaluation of the positional precision of chromatin loops

We used bioframe.overlap function (v0.3.3) to intersect loop anchors with CTCF and RAD21 (ChIP–seq/CUT&RUN) peaks, as well as ATAC-seq peaks [29]. These overlapping percentages are shown in Figs. 4c, 6b, S5c, S8b, and S9.

### Visualization and quantification of chromatin loops

#### Loop pileups

The pileups were generated by averaging stacks of individual loop matrices produced by the cooltools.pileup function at two resolutions, 5 kb with 50 kb flanking regions and 10 kb with 100 kb flanking regions.

#### Upset plots

We generated UpSet plots to visualize the number of detected loops shared among one or more samples, using the ComplexHeatmap R package (v2.10.0) [30].

#### Stackup plots

We created stackup plots of individual loops by extracting the middle rows (flanked upstream anchors) from loop matrices generated by the cooltools.pileup function (v0.5.2) and sorting them according to the average signal in the middle four bins.

#### Loop strength

Using the cooltools.pileup function, 5 kb or 10 kb resolution interaction matrices of 21 × 21 bins were first generated, centered on each loop. Loop strength was then calculated as the mean signal within the central 3 × 3 bin square divided by the mean signal from three neighboring regions (upper left, upper right, and lower right), following the approach of a previous study (Additional file 1: Fig. S7) [15]. These quantifications are shown in Figs. 4b, 5a, S5b, S6a–b, S11, and S10.

### Detection of gained and lost loops between two samples using DESeq2

We first compiled the UnionLoops-defined lost and gained loops for subsequent pairwise comparison by DESeq2: 5 kb loops from 0 h to 72 h for the Bond et al. dataset, and 10 kb loops from the DE to PP stages for the Lyu et al. dataset. We then extracted the unnormalized contact count matrix for these two loop sets from the .mcool files of their biological replicates (four for the Bond et al. dataset and two for the Lyu et al. dataset) using the cooler.Cooler.matrix() (v0.9.3) selector with balance=False, thereby omitting ICE normalization and retaining raw counts for subsequent differential analysis. The resulting count matrix was analyzed with DESeq2 (v1.34.0) using the design formula ∼ rep + time, which accounts for replicate-specific variation (rep) while testing for differences associated with the experimental time factor (time) [17]. Loops were classified as gained if they had a logL fold change > 0 with an adjusted P-value < 0.05, and as lost if they had a logL fold change < 0 with an adjusted P-value < 0.05.

### Fine-tuning of HiCCUPS loops via Mariner-style, intermediate, and UnionLoops strategies

To test the influence of two operational features on fine-tuning shown in Additional file 1: Fig. S8a, we used the HiCCUPS loops as a reference (i.e., same loci) to compare the improvement of their positional precision using Mariner-style, intermediate, and UnionLoops strategies. As shown in Additional file 1: Fig. S8b, the Mariner-style strategy clustered only the final filtered HiCCUPS loops from individual samples and selected the cluster representative as the pixel with the maximum value in any single sample [13]. The Intermediate strategy also clustered the final filtered HiCCUPS loops but identified the representative pixel as the one with the maximum sum across all samples in the cluster. In contrast, UnionLoops clustered pre-filtered significant pixels (e.g., enriched pixels from HiCCUPS) across samples and identified the representative pixel as the one with the maximum sum across all samples. To control for confounding factors, all three strategies employed the same clustering method as used in HiCCUPS, with a cluster radius of 20 kb, and all used normalized contacts as the metric for representative selection.

### Fine-tuning of SIP loops via Mariner and UnionLoops

As in Bond et al., we applied the mergePairs function in the Mariner R package (v1.6.0) to cluster 5 kb SIP loops at 0 h, 6 h, 72 h, and MEGA from the K562 Hi-C datasets [13]. Clusters were assigned to representative pixels with the maximum APScoreAvg value (a SIP loop caller output column) among the four samples, using the parameters radius = 10000, column = “APScoreAvg”, and selectMax = TRUE. We upgraded the UnionLoops nextflow pipeline from v1.0.0 to v1.1.0 to support merging loops from multiple datasets generated by external loop callers, enabling integration of these loop sets alongside built-in HiCCUPS calls. Using UnionLoops v1.1.0, we clustered 5 kb SIP-derived loops from individual 0 h, 6 h, 72 h, and MEGA Hi-C maps using the same HiCCUPS clustering method applied to enriched pixels. Within each cluster, we selected a representative pixel defined as the locus with the maximum sum of normalized contact across all datasets. This procedure is identical to the built-in HiCCUPS workflow in UnionLoops (Fig. 2), except that the input comprises loop pixels derived from external loop callers. No additional filtering was applied, as these loop callers already implement their own statistical thresholds and the HiCCUPS-specific thresholds are not applicable to these external loops. The analysis was performed using the parameters assembly_name = “hg38”, dots_clustering_radius = 20000, resolution = 5000, and flank = 50000.

### Fine-tuning of Mustache loops via UnionLoops

We first used Mustache (v1.3.3) with the parameters ‘-r 5000 -pt 0.05’ to separately generate loop sets at 5kb resolution using 0 h, 6 h, 72 h, and MEGA mcool files from the K562 Hi-C datasets [16]. Using the same approach as for fine-tuning SIP loops, we applied UnionLoops (v1.1.0) to merge 5 kb Mustache-derived loop calls from individual 0 h, 6 h, 72 h, and MEGA Hi-C maps, generating a union loop set across time points. The analysis was conducted with the parameters assembly_name = “hg38”, dots_clustering_radius = 20000, resolution = 5000, and flank = 50000.

## Declarations

### Peer Review Information

Daan Noordermeer and Wenjing She were the primary editors of this article and managed its editorial process and peer review in collaboration with the rest of the editorial team. The peer-review history is available in the online version of this article.

### Ethics Approval and Consent to Participate

Not applicable

### Consent for Publication

Not applicable

### Availability of Data and Materials

The UnionLoops nextflow pipeline (unionloops-nf) is available at https://github.com/dekkerlab/unionloops-nf under the MIT License and is archived on Zenodo at DOI: https://doi.org/10.5281/zenodo.19837477 [31,32]. All scripts and Jupyter Notebooks for the UnionLoops manuscript are available through https://github.com/dekkerlab/unionloops_paper under the MIT licence, and deposited in Zenodo via the following DOI: https://doi.org/10.5281/zenodo.19837282 [33,34].

Hi-C data (K562_0h, K562_6h, K562_72h) were generated in D. Phanstiel’s laboratory and can be found in the NCBI Gene Expression Omnibus (GEO; https://www.ncbi.nlm.nih.gov/geo/) under accession number GSE214123 [35]. Hi-C data (hESC, DE, PGT, PP, SC-β organoids) were generated in V. Corces’s laboratory and are available in the GEO under accession number GSE210524 [36]. For Hi-C data (K562_0h, K562_6h, K562_72h), we also downloaded SIP loops at 5kb resolution for each time point, as well as the MEGA file under accession number GSE214123 [35].

We downloaded 1) processed bigwig and peak lists for CUT&RUN data of CTCF and RAD21 under accession number GSE213908; 2) processed peak lists for ATAC-seq data under accession number GSE213295 for the study of K562 differentiation [37,38]. For the study of pancreatic cell differentiation, processed peak lists for ChIP-seq data of CTCF and RAD21 were downloaded under accession number GSE211101 [39]. We downloaded raw FASTQ files of ATAC-seq data from H9 hESC under accession number GSE85330, generated in M. Snyder’s laboratory [40]. We also obtained raw FASTQ files of ATAC-seq data of DE, PGT, PP, and SC-β organoids under accession number GSE211101 [39].

### Authors’ Contributions

J.L. performed all analyses. All authors conceived of the project and wrote the manuscript. All authors read and approved the final manuscript.

## Acknowledgements

We thank members of the Dekker lab for discussions and advice. We thank Dr. Caryn Navarro for editing the manuscript. We thank Drs. Eileen Furlong and Mattia Forneris for suggesting Mustache in conjunction with UnionLoops.

## Supplementary Information

### Additional file 1: PDF document containing all supplementary figures

**Additional file 1: Fig. S1:** UnionLoops corrects false dataset specificity of chromatin loops in the conventional method, as demonstrated by biological replicate comparisons.

**Additional file 1: Fig. S2:** Schematic plot illustrating UnionLoops’ additional filtering of singletons whose anchors do not overlap with boundaries of any non-singleton clusters after pooling enriched pixels from all samples.

**Additional file 1: Fig. S3:** Percentage of loops rescued exclusively by MEGA among all rescued loops at each time point.

**Additional file 1: Fig. S4:** In contrast to centroids, a substantial number of singletons in each sample remained unclustered and lacked adjacent enriched pixels after pooling across all samples.

**Additional file 1: Fig. S5:** UnionLoops improves sensitivity and positional precision of HiCCUPS loop calls at five time points.

**Additional file 1: Fig. S6:** UnionLoops improves loop specificity across hESC, DE, PGT, PP, and SC-β organoids compared to the conventional method.

**Additional file 1: Fig. S7:** Quantification of loop strength based on local enrichment.

**Additional file 1: Fig. S8:** The differences in two operational features allow UnionLoops to uniquely rescue filtered loops and achieve better positional precision than Mariner.

**Additional file 1: Fig. S9:** UnionLoops improves loop precision for SIP and Mustache loop callers.

**Additional file 1: Fig. S10:** UnionLoops improves loop specificity for both SIP and Mustache loop callers compared to the conventional method.

**Additional file 1: Fig. S11:** Loop strength trajectories of loops shared across all samples.

### Additional file 2: PDF document containing all supplementary tables

**Additional file 2: Table S1:** Number of chromatin loops per sample detected by HiCCUPS and UnionLoops in each dataset at different resolutions.

**Additional file 2: Table S2:** Summary of differential chromatin loops detected by DESeq2 using UnionLoops-defined lost and gained loop sets.

