## Additional File 1 for "UnionLoops: a workflow for calling chromatin loops across related Hi-C datasets with improved specificity, precision, and sensitivity"

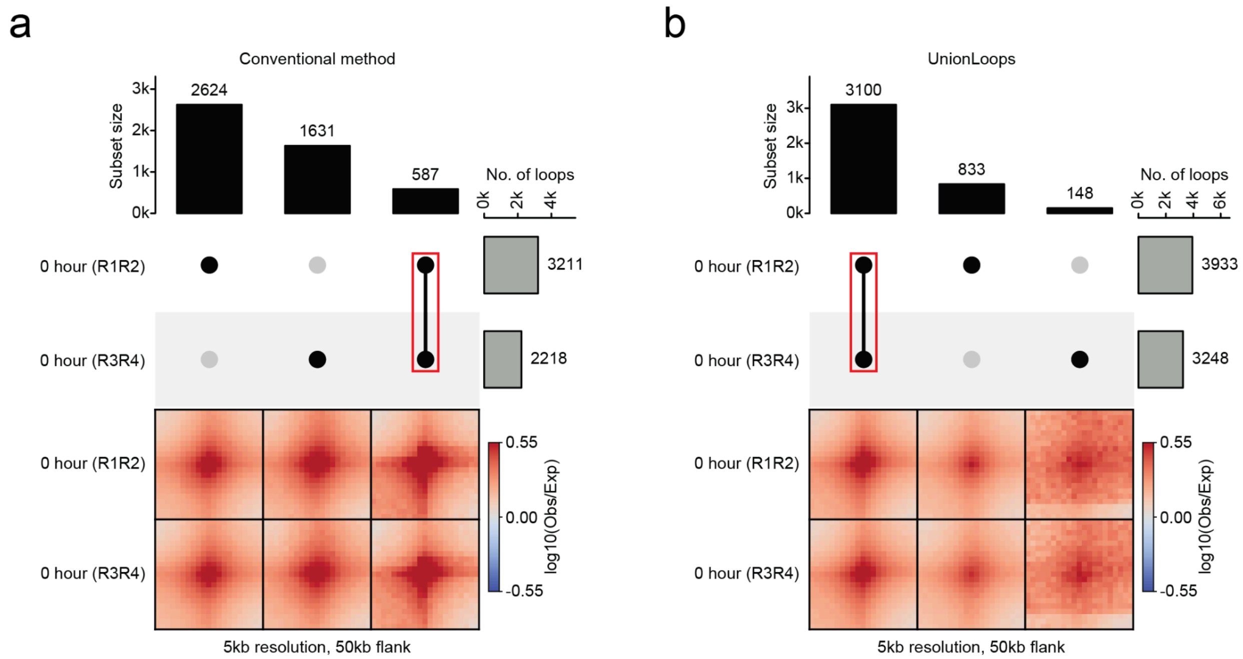


**Fig. S1. UnionLoops corrects false dataset specificity of chromatin loops in the conventional method, as demonstrated by biological replicate comparisons. a**, UpSet plots (top) show chromatin loops detected by the conventional method across two merged Hi-C datasets from replicates R1–R2 and R3–R4 at 0 hour (Bond et al.), revealing a large number of dataset-specific loops. Red rectangles highlight loops detected in both datasets. Pileups (bottom) display averaged contact signals for each loop category defined above. **b**, Same as in **a**, but for UnionLoops, which recovers predominantly shared loops between biological replicates at 0 hours.

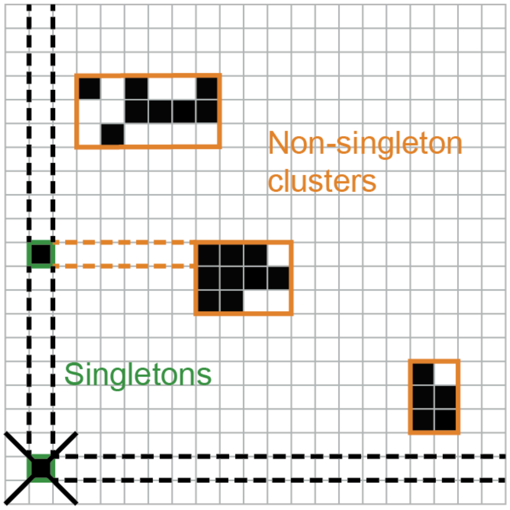


**Fig. S2. Schematic plot illustrating UnionLoops' additional filtering of singletons whose anchors do not overlap with boundaries of any non-singleton clusters after pooling enriched pixels from all samples.**

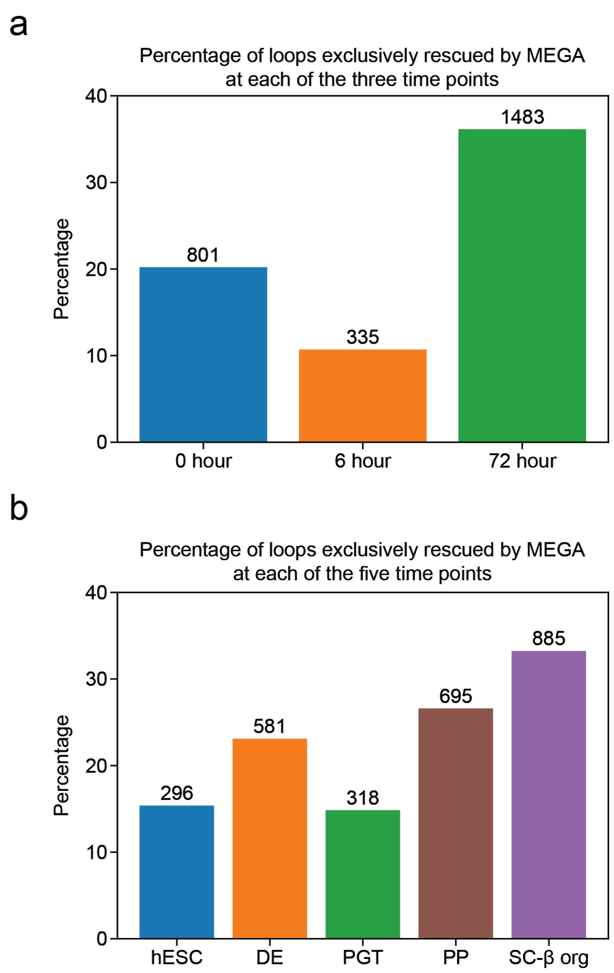


**Fig. S3. Percentage of loops rescued exclusively by MEGA among all rescued loops at each time point.** For each time point, the proportion was calculated by dividing the number of loops rescued exclusively by MEGA by the total number of rescued loops. The labels above the bars show the corresponding numbers of loops. **a**, three-timepoint in situ Hi-C data from Bond et al. **b**, five-timepoint in situ Hi-C data from Lyu et al.

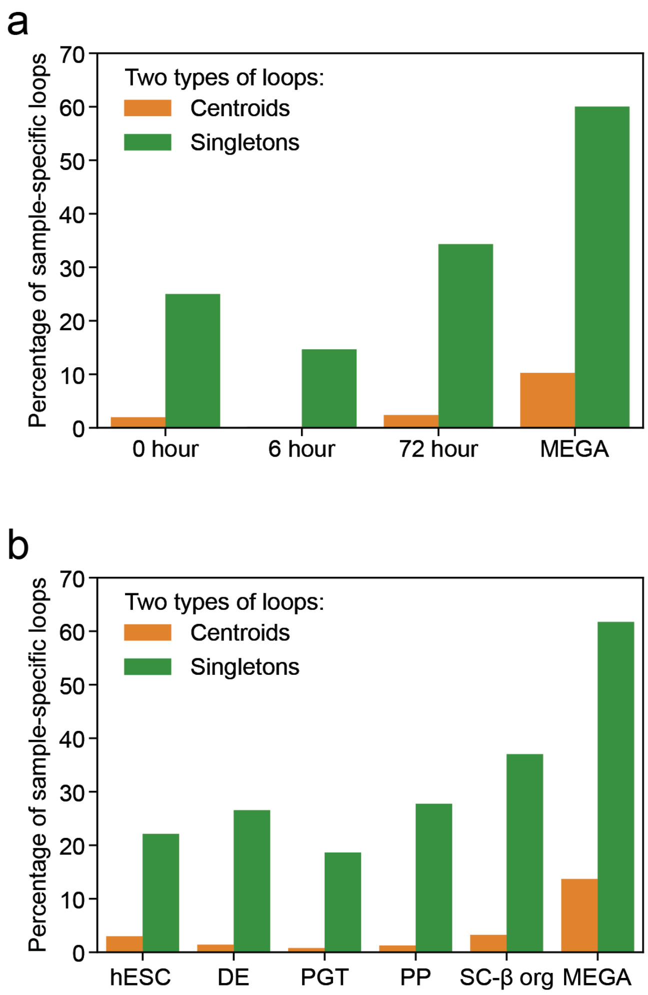


**Fig. S4. In contrast to centroids, a substantial number of singletons in each sample remained unclustered and lacked adjacent enriched pixels after pooling across all samples.** For each sample, the proportion was calculated by dividing the number of sample-specific loops by the total number of loops. **a**, Three-timepoint in situ Hi-C data from Bond et al. **b**, Five-timepoint in situ Hi-C data from Lyu et al.

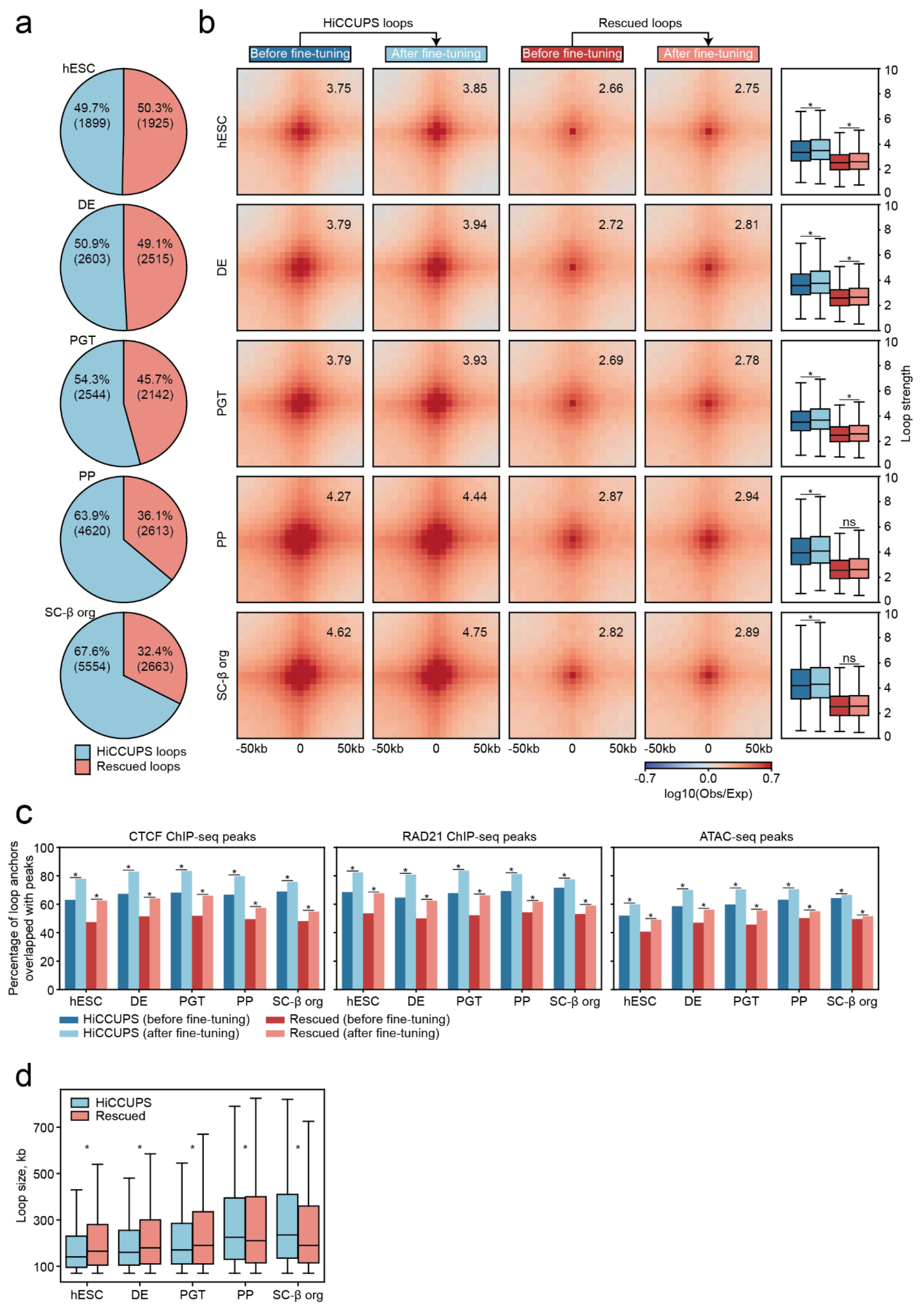


**Fig. S5. UnionLoops improves sensitivity and positional precision of HiCCUPS loop calls at five time points. a**, The percentage of HiCCUPS loops versus the percentage of rescued loops. **b**, Pileups of HiCCUPS and rescued loops before and after fine-tuning, with corresponding loop strength distributions. The averaged loop strength value in each pileup represents signal enrichment relative to the local background (see Methods). Wilcoxon rank-sum test; asterisks indicate $P < 0.05$. **c**, The percentages of anchors from HiCCUPS and rescued loops, before and after fine-tuning, that overlapped with CTCF ChIP-seq, RAD21 ChIP-seq, and ATAC-seq peaks. Chi-square test of independence; asterisks indicate $P < 0.05$. **d**, Loop size comparisons between HiCCUPS and rescued loops. Wilcoxon rank-sum test; asterisks indicate $P < 0.05$. Loop resolution is 5 kb for all panels (**a–d**).

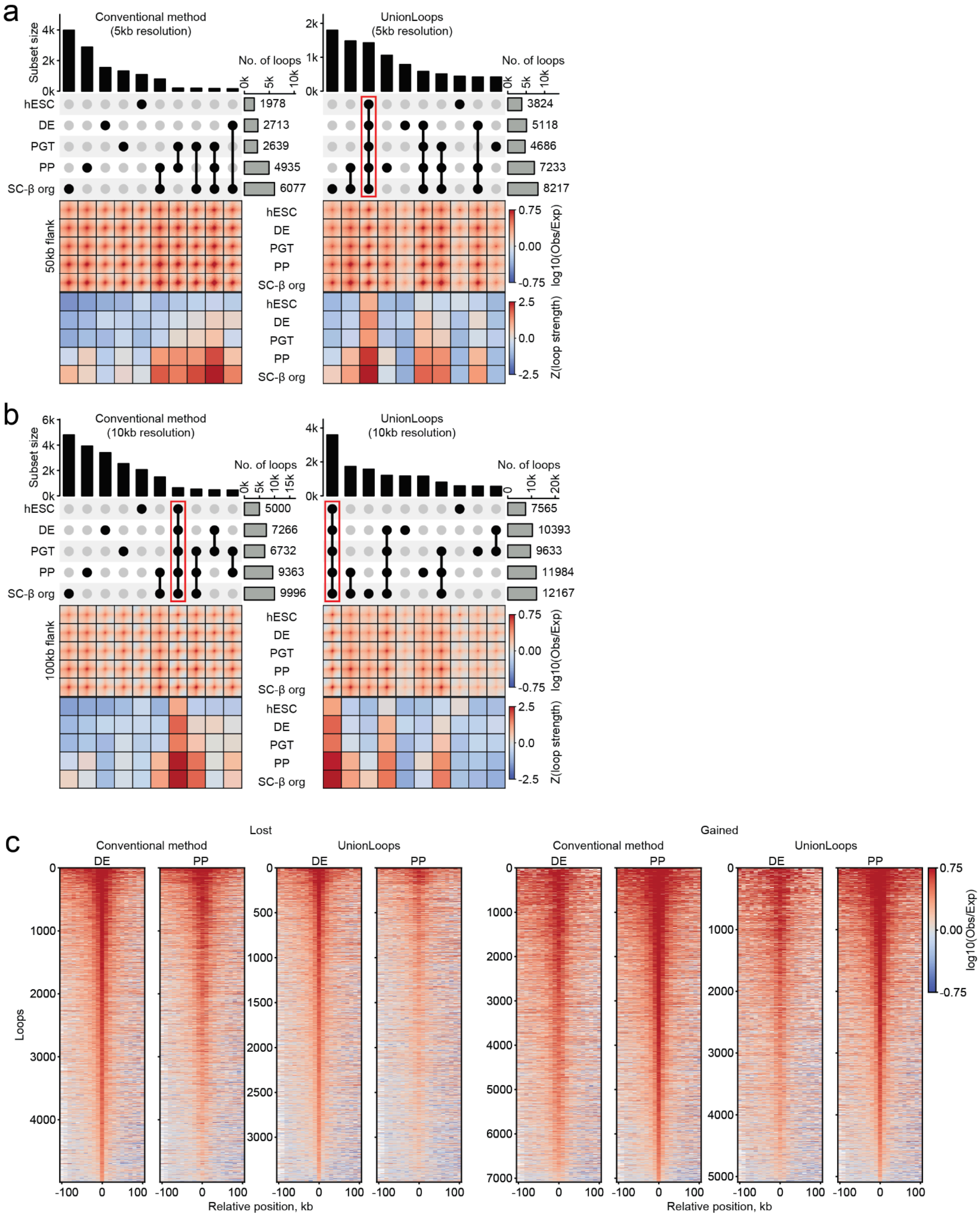


**Fig. S6. UnionLoops improves loop specificity across hESC, DE, PGT, PP, and SC-β organoids compared to the conventional method. a-b**, Combined visualization of loop detection and signal strength across five time points at 5 kb (**a**) and 10 kb (**b**) resolutions. Each panel includes UpSet plots (top), pileups (middle), and heatmaps of loop strength matrices (bottom) for the top 10 most frequent loop subsets, as shown in the same way as in **Fig. 5a. c**, Stackup plots at 100kb-flanked upstream 10 kb anchors of individual lost and gained loops between DE and PP.

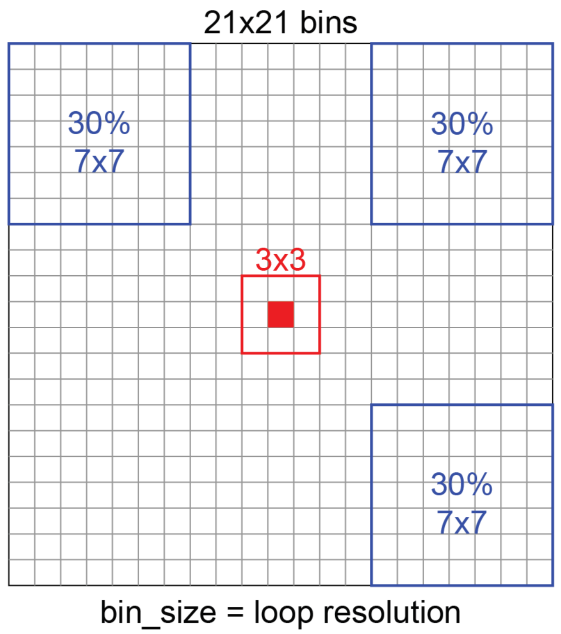


**Fig. S7. Quantification of loop strength based on local enrichment.** Loop strength was measured by calculating enrichment relative to the local background. A 21 × 21 observed-over-expected contact matrix was generated for each loop, centered on the loop position and using the loop resolution as the bin size. Loop strength was defined as the ratio between the average signal in the central 3 × 3 region and the average of three neighboring regions: upper left, upper right, and lower right.

a

| Operational features | Mariner | UnionLoops |
|---|---|---|
| Input pixels for clustering | Final loops (**FL**) after additional filtering | Significant pixels (**SP**) before additional filtering |
| Representative per cluster | Pixel with the maximum value (**MV**) in one of the involved samples | Pixel with the maximum sum (**MS**) across all involved samples |

b

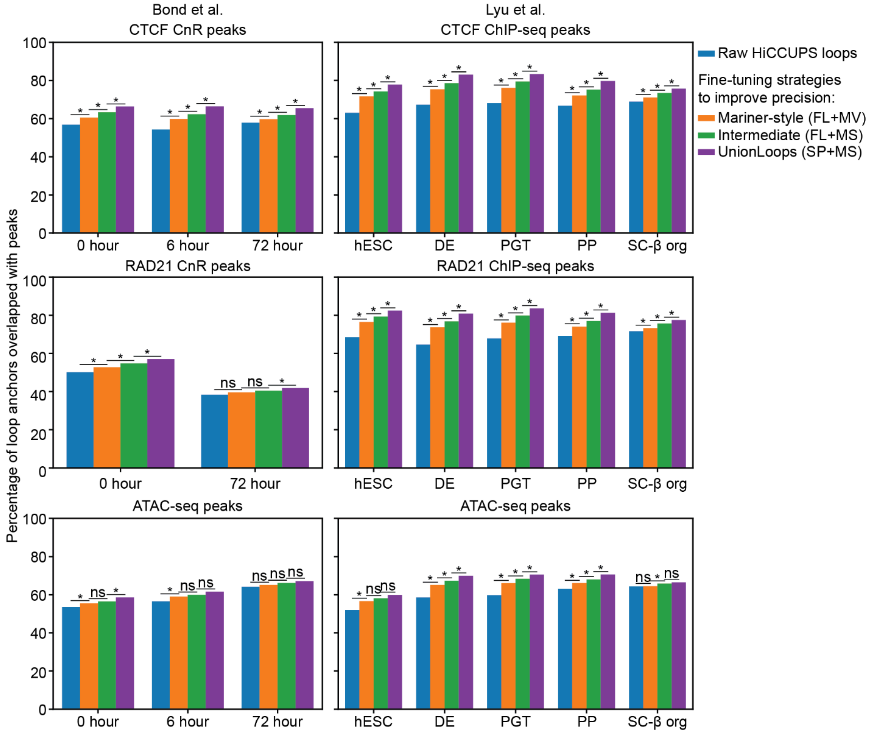


c

| Output features | Mariner | UnionLoops |
|---|---|---|
| Rescued loops | ✗ (FL) | ✓ (SP) |
| Improved precision | ✓ (FL+MV) | ✓✓ (SP+MS) |

**Fig. S8. The differences in two operational features allow UnionLoops to uniquely rescue filtered loops and achieve better positional precision than Mariner. a**, The table compares two key operational features of Mariner and UnionLoops: the input pixels for clustering and the method for defining a representative per cluster. **b**, The percentages of anchors overlapping with CTCF, RAD21 (CnR/ChIP-seq), and ATAC-seq peaks for both the Bond et al. and Lyu et al. datasets are shown for raw HiCCUPS loops and three fine-tuned loop sets generated using strategies based on two operational features: Mariner-style (FL+MV), Intermediate (FL+MS), and UnionLoops (SP+MS) (see Methods). Chi-square test of independence; asterisks indicate P < 0.05. **c**, The table summarizes differences in two output features (rescued loops and improved positional precision) and their associated operational features between Mariner and UnionLoops. Codes of operational features used in all panels (**a-c**): FL (final loops), SP (significant pixels), MV (maximum value), and MS (maximum sum). Loop resolution: 5 kb.

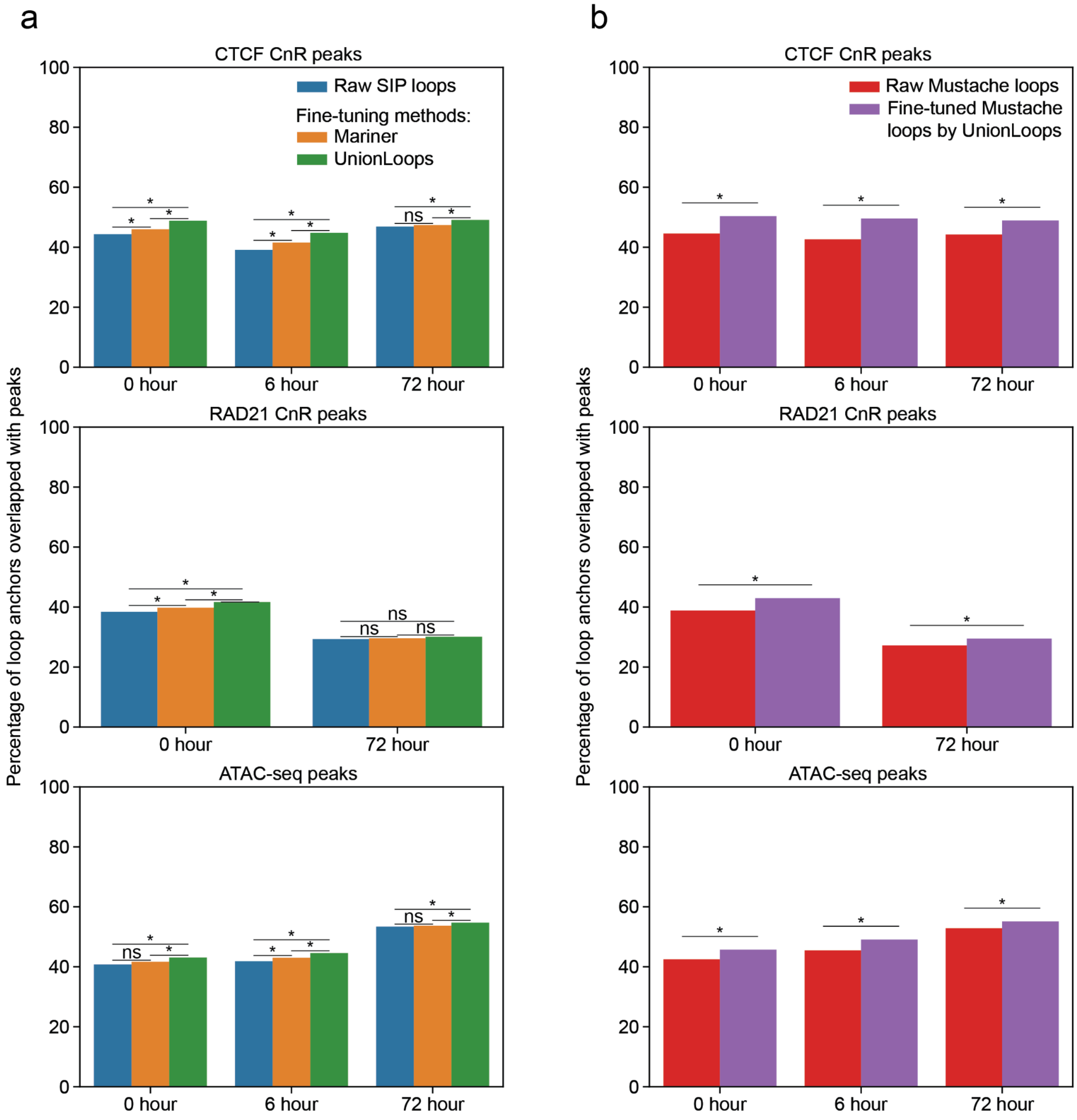


**Fig. S9. UnionLoops improves loop precision for SIP and Mustache loop callers. a-b,** The percentages of anchors overlapping with CTCF, RAD21 (CnR), and ATAC-seq peaks in the Bond et al. dataset are shown for (**a**) raw SIP loops and their fine-tuned loop sets using Mariner and UnionLoops and (**b**) raw Mustache loops and their fine-tuned loop sets using UnionLoops (see Methods), as shown in the same way as in **Figs. 4c and S5c.** Chi-square test of independence; asterisks indicate $P < 0.05$. Loop resolution: 5 kb.

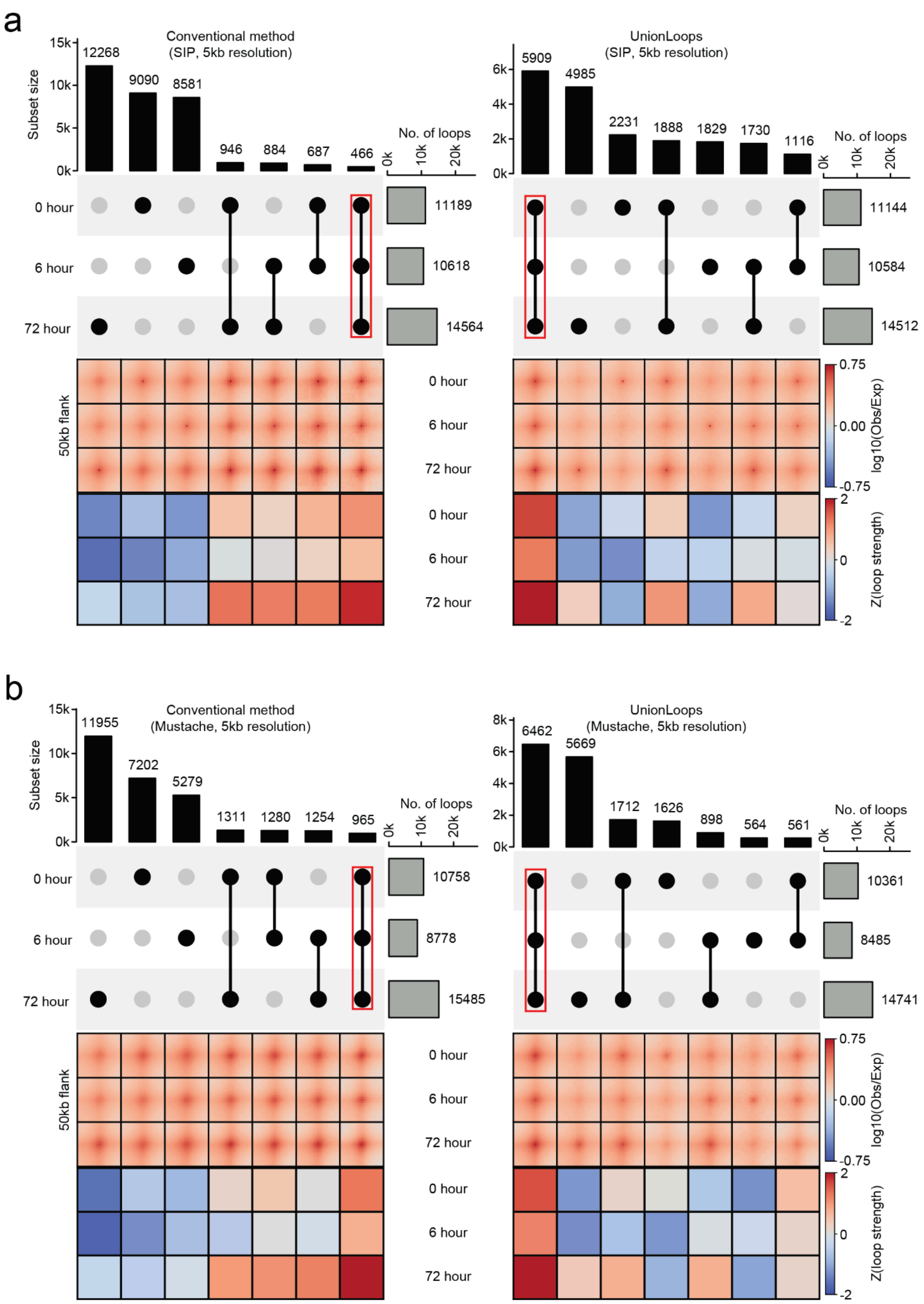


**Fig. S10. UnionLoops improves loop specificity for both SIP and Mustache loop callers compared to the conventional method. a-b,** Combined visualization of loop detection and signal strength across the three time points for loops at 5kb resolution generated from (**a**) SIP and (**b**) Mustache loop callers, as shown in the same way as in **Figs. 5a and S6a-b**.

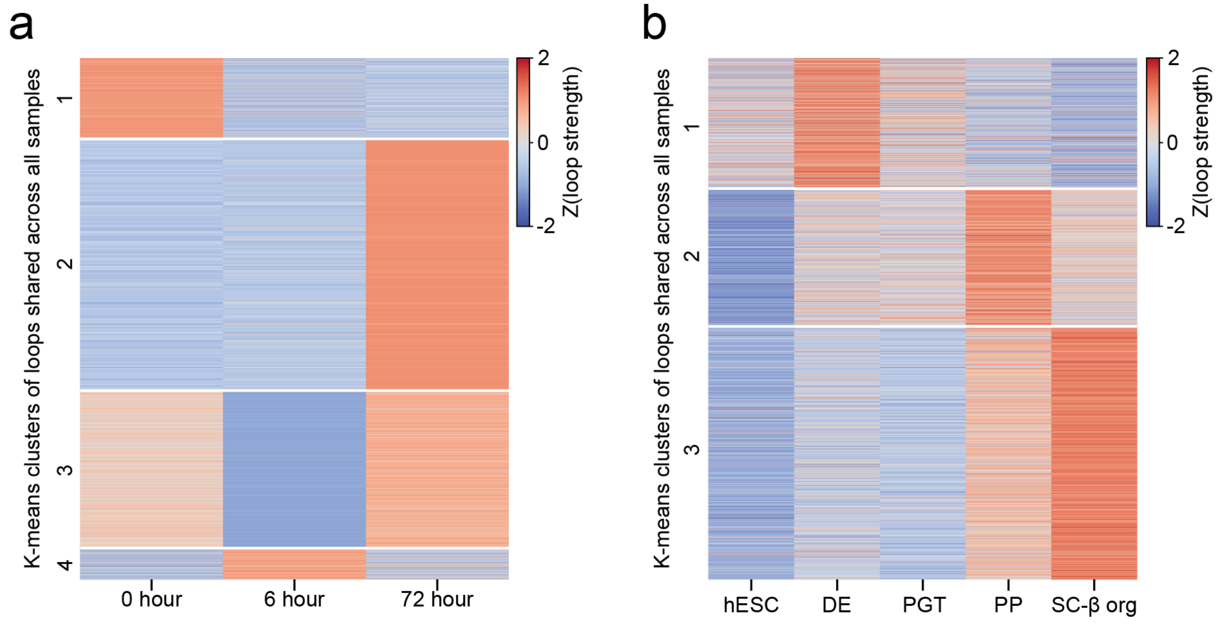


**Fig. S11. Loop strength trajectories of loops shared across all samples.** Loop strength was tracked across samples for loops identified in all samples by UnionLoops. Values were Z-score normalized per loop across samples before K-means clustering. Each trajectory represents dynamic changes in loop strength over time. **a**, three-timepoint in situ Hi-C data from Bond et al. (5 kb resolution). **b**, five-timepoint in situ Hi-C data from Lyu et al. (10 kb resolution).
