## Additional File 2 for "UnionLoops: a workflow for calling chromatin loops across related Hi-C datasets with improved specificity, precision, and sensitivity"

| Dataset | Resolution | Sample | HiCCUPS | UnionLoops |
|---|---|---|---|---|
| [35] | 5 kb | 0 hour | 4088 | 7791 |
| | | 6 hour | 3030 | 5979 |
| | | 72 hour | 5818 | 9415 |
| | | MEGA | 5319 | 12271 |
| [36] | 5 kb | hESC | 1978 | 3824 |
| | | DE | 2713 | 5118 |
| | | PGT | 2639 | 4686 |
| | | PP | 4935 | 7233 |
| | | SC-β org | 6077 | 8217 |
| | | MEGA | 12866 | 13351 |
| [36] | 10 kb | hESC | 5000 | 7565 |
| | | DE | 7266 | 10393 |
| | | PGT | 6732 | 9633 |
| | | PP | 9363 | 11984 |
| | | SC-β org | 9996 | 12167 |
| | | MEGA | 16351 | 19104 |

**Table S1. Number of chromatin loops per sample detected by HiCCUPS and UnionLoops in each dataset at different resolutions.**

| Dataset | UnionLoops-defined | Total | Lost (DESeq2) | Gained (DESeq2) | Non-significant (DESeq2) |
|---|---|---|---|---|---|
| [35] (0 h to 72 h; 5 kb; 4 reps) | Lost | 1744 | 28 | 0 | 1716 |
| | Gained | 3368 | 0 | 11 | 3357 |
| [36] (DE to PP; 10 kb; 2 reps) | Lost | 3500 | 0 | 0 | 3500 |
| | Gained | 5091 | 0 | 0 | 5091 |

**Table S2. Summary of differential chromatin loops detected by DESeq2 using UnionLoops-defined lost and gained loop sets.**
